# The pseudoknot structure of a viral RNA reveals a conserved mechanism for programmed exoribonuclease resistance

**DOI:** 10.1101/2024.12.17.628992

**Authors:** Jeanine G. Gezelle, Sophie M. Korn, Jayden T. McDonald, Zhen Gong, Anna Erickson, Chih-Hung Huang, Feiyue Yang, Matt Cronin, Yen-Wen Kuo, Brian T. Wimberly, Anna-Lena Steckelberg

**Affiliations:** Department of Biochemistry and Molecular Biophysics, Columbia University, New York, NY, USA; Department of Plant Pathology, University of California, Davis, CA, USA; Department of Systems Biology, Columbia University, New York, NY, USA; New York Structural Biology Center, New York, NY, USA

**Author notes:** Corresponding author: Contact: Anna-Lena Steckelberg.

## Abstract

Exoribonuclease-resistant RNAs (xrRNAs) are viral RNA structures that block degradation by cellular 5′-3′ exoribonucleases to produce subgenomic viral RNAs during infection. Initially discovered in flaviviruses, xrRNAs have since been identified in wide range of RNA viruses, including those that infect plants. High sequence variability among viral xrRNAs raises questions about the shared molecular features that characterize this functional RNA class. Here, we present the first structure of a plant-virus xrRNA in its active exoribonuclease-resistant conformation. The xrRNA forms a 9 base pair pseudoknot that creates a knot-like topology similar to that of flavivirus xrRNAs, despite lacking sequence similarity. Biophysical assays confirm a compact pseudoknot structure in solution, and functional studies validate its relevance both *in vitro* and during infection. Our study reveals how viral RNAs achieve a common functional outcome through highly divergent sequences and identifies the knot-like topology as a defining feature of xrRNAs.

---

Positive-sense single-stranded RNA (+RNA) viruses include many important human pathogens that pose major global health threats^1,2^. Through evolution, these obligate intracellular parasites have optimized their compact genomes and developed sophisticated strategies to hijack host cell machinery, often using highly structured genomic RNA elements to regulate gene expression and evade immune responses^3,4^. Viral RNA structures hold great potential as antiviral targets, but their molecular mechanisms remain largely unexplored^5–9^. A notable example are the exoribonuclease-resistant RNA (xrRNA) elements, which block degradation by cellular 5′-3′ exoribonucleases, including the highly processive cytoplasmic enzyme Xrn1, at specific sites in the viral genome^10–13^. This mode of programmed exoribonuclease resistance effectively transforms cellular exoribonucleases into RNA maturation enzymes, trimming viral genomes to produce subgenomic RNAs (sgRNAs) (Extended Data Fig. 1a). The mechanism is particularly remarkable because Xrn1 and related enzymes typically degrade even highly structured RNA substrates rapidly and without releasing degradation intermediates, a process facilitated by the enzymes’ translocation-coupled RNA-unwinding activity^14–16^.

Viral xrRNAs were discovered over a decade ago in West Nile virus (WNV)^10^, and have since been identified in the genomes of many related mosquito-borne flaviviruses (MBFV), including the human-pathogenic Dengue virus (DENV), Japanese encephalitis virus (JEV), Yellow fever virus (YFV) and Zika virus (ZIKV), as well as in all other genera of the *Flaviviridae* family (i.e., in Pegiviruses, Pestiviruses and Hepaciviruses)^17–19^. Flaviviruses have +RNA genomes with a single open reading frame (ORF) flanked by highly structured 5′ and 3′ untranslated regions (UTRs)^20^. XrRNAs are located at the start of the flaviviral 3′ UTR, forming a compact double-pseudoknot (PK) structure centered around a 3-way junction (3WJ) that creates a protective ring around the 5′ end of the RNA element^11,21,22^. Characterization of conserved sequence and structural features of flaviviral xrRNAs lead to a subdivision into class 1a, class 1b and class 2 xrRNAs (Extended Data Fig. 1b)^13,17,18,23^. In all classes, the RNA adopts a slipknot-like topology that provides directional stability; the xrRNA acts as a molecular brace, blocking exoribonucleases from the 5′ end while allowing unwinding by the viral RNA-dependent RNA polymerase (RdRP) approaching from the 3′ end^11,12,22–27^. Stalling of Xrn1 in the flaviviral 3′ UTR leads to the production of non-coding subgenomic flavivirus RNAs (sfRNAs), which play important roles in immune modulation and viral pathogenicity^10,28–38^. Although the exact biological functions of sfRNAs are not fully understood, disrupting their production results in viral attenuation, and xrRNAs have emerged as promising targets for antiviral therapies and live-attenuated vaccines^39^.

Initially considered an idiosyncrasy of flaviviruses, xrRNAs have now been identified in many plant-infecting +RNA viruses from the *Solemoviridae* and *Tombusviridae* families, where they generate both non-coding and protein-coding sgRNAs^40–42^. This discovery has established RNA structure-based exoribonuclease-resistance as a widespread viral mechanism for sgRNA production. Surprisingly, members of the new xrRNA class are significantly smaller and show no sequence similarity to flavivirus xrRNAs^13,40,42^ (Extended Data Fig. 1b). This disparity reveals that diverse RNA sequences can effectively block exoribonucleases and raises intriguing questions about the shared molecular features that define this emerging class of functional RNAs. Following the naming convention for flavivirus xrRNAs, we classify these new exoribonuclease-resistant viral RNA structures as class 3 xrRNAs^13^ (Extended Data Fig. 1b). In contrast to the double-pseudoknot structure of flaviviral class 1 and class 2 xrRNAs, class 3 xrRNA contain a single stem loop (SL) formed by coaxially stacked helices P1 and P2, with the potential to form a PK between the apical loop and the 3′ end of the RNA (Extended Data Fig. 1b)^40,42^. As in the flaviviral xrRNA, PK formation creates an exoribonuclease-resistant structure. However, the PK is not always formed^40,42^. Instead, class 3 xrRNAs exist as a dynamic conformational ensemble, switching between open (SL) and closed (PK) structures^40^. Single-molecule Förster resonance energy transfer measurements and site-directed mutagenesis experiments revealed a coordinated co-degradational refolding mechanism, in which partial unwinding of P1 by Xrn1 shifts the structural equilibrium to stabilize the exoribonuclease-resistant PK^40^. Previously solved high-resolution crystal structures of class 3 xrRNAs – xrRNAs from sweet clover necrotic mosaic virus, (SCNMV, *Tombusviridae)*^40^ and potato leaf roll virus (PLRV, *Solemoviridae*)^42^ – captured the RNA in its open SL conformation, representing the pre-degradation folding intermediate. In both structures, the PK interactions are formed *in trans* as part of a crystallographic dimer of two adjacent RNA molecules. Modelling the PK conformation *in cis* suggested that it would create a similar protective ring topology as flaviviral xrRNAs^40^. However, without a high-resolution structure of a class 3 xrRNA with the intramolecular PK formed, the full set of molecular interactions stabilizing the PK conformation remained unknown.

In this study, we combine structural biology, biophysics and molecular virology to provide the most detailed mechanistic insights into the class 3 xrRNA to date, including the first high-resolution crystal structure of a class 3 xrRNA in its active PK conformation. We focus on an xrRNA from the 3′ UTR of beet western yellows virus ST9-associated RNA (referred to as ST9-associated, ST9a), a subviral +RNA that replicates autonomously but requires a co-infecting polerovirus for encapsidation and transmission^43^. Our structure reveals that the ST9a xrRNA forms a 9 base pair PK and additional long-range interactions to adopt the characteristic protective ring topology. We use biophysical assays to confirm that the compact PK structure forms in solution, and validate the functional relevance of the interactions using site-directed mutagenesis both *in vitro* and during viral infection. Structural features of the ST9a xrRNA closely resemble previously solved open xrRNA structures from PLRV and SCNMV, suggesting that this structure can serve as a model for the PK conformation of the entire xrRNA class. Most importantly, the structure allows us to directly compare the molecular architecture of two highly divergent xrRNA classes: flavivirus xrRNA and plant-virus xrRNAs. We confirm expected similarities, such as the presence of a protective ring, but also uncover unexpected parallels between these classes, pointing to common features that may be universally shared by viral xrRNAs.

## Results

### A compact xrRNA exists in the 3′ UTR of the ST9a replicon

The xrRNAs identified so far likely represent only a fraction of the overall diversity, with many more to be discovered. In line with this, several new putative xrRNA structures have recently been found in viral RNA genomes, though their detailed structures remain unknown. One such newly identified putative xrRNA is located in the 3′ UTR of ST9a^44^. Because ST9a requires a co-infecting plant virus for transmission^45,46^, this discovery represents the first instance of an xrRNA identified outside the genome of a fully autonomous RNA virus. A previous study demonstrated that ST9a produces sgRNAs during infection and that a sequence in the viral 3′ UTR resists degradation by recombinant Xrn1 *in vitro*, but the underlying structural mechanisms were not explored^44^. We used agroinfection of *Nicotiana benthamiana* (*N. benthamiana*) plants to confirm the production of sgRNAs during ST9a infection. Northern blotting with a probe against the 3′ UTR revealed the presence of a ∼400 nucleotide-long sgRNA that is coterminal with the ST9a genome (Fig. 1a, Extended Data Fig. 2). A less abundant, faster-migrating RNA species was also detected by northern blotting. This band might be caused by sgRNA processing in host cells, as has been previously demonstrated for ZIKV sfRNAs^47^, or by host ribosomal RNA (rRNA) trapping^48^. To verify the presence of an authentic xrRNA element capable of blocking exoribonuclease degradation independently of *trans*-acting proteins, we challenged *in vitro*-transcribed, purified viral 3′ UTR sequences from ST9a with recombinant yeast Xrn1, a close homolog of the plant 5′-3′ exoribonuclease Xrn4^16^ (Extended Data Fig. 3a-b). An Xrn1-resistant degradation product was detected within minutes of incubation and remained stable for several hours in the presence of Xrn1 (Extended Data Fig. 3c), demonstrating that the ST9a 3′ UTR contains a highly efficient nuclease-resistant RNA element. To map the Xrn1 stop site, we reverse-transcribed the degradation products and analyzed the cDNA via sequencing gel electrophoresis (Fig. 1b). Notably, the 5′ end of the resistant fragments mapped to two nucleotides in the 3′ UTR that precisely matched the previously determined 5′ end of sgRNAs produced during ST9a infection^44^, confirming that our *in vitro* assay faithfully recapitulates exoribonuclease resistance observed *in vivo*. To identify the minimal RNA element required for exoribonuclease-resistance, we treated 3′-truncated RNA substrates with Xrn1 and found that a 54-nucleotide segment was necessary and sufficient to confer full resistance (Fig. 1c). Closer inspection of the minimal xrRNA sequence revealed similarities to the previously identified class 3 xrRNAs^40–42^. Like all members of the xrRNA class 3, the ST9a xrRNA has the potential to form a short SL with stacked helices P1 and P2, along with a PK between the apical loop and the 3′ end (Fig. 1d). However, we found notable differences between ST9a xrRNA and other class 3 xrRNA members that likely explain why its sequence was not previously identified in bioinformatic searches for related structures^41^. Most importantly, ST9a xrRNA has the potential to form a significantly extended PK–nearly twice as long as those observed in other class 3 xrRNAs (Fig. 1d)^40–42^. This unique tertiary interaction likely stabilizes the xrRNA in its closed PK conformation, making the ST9a xrRNA an excellent candidate for structural studies.

**Figure 1.**
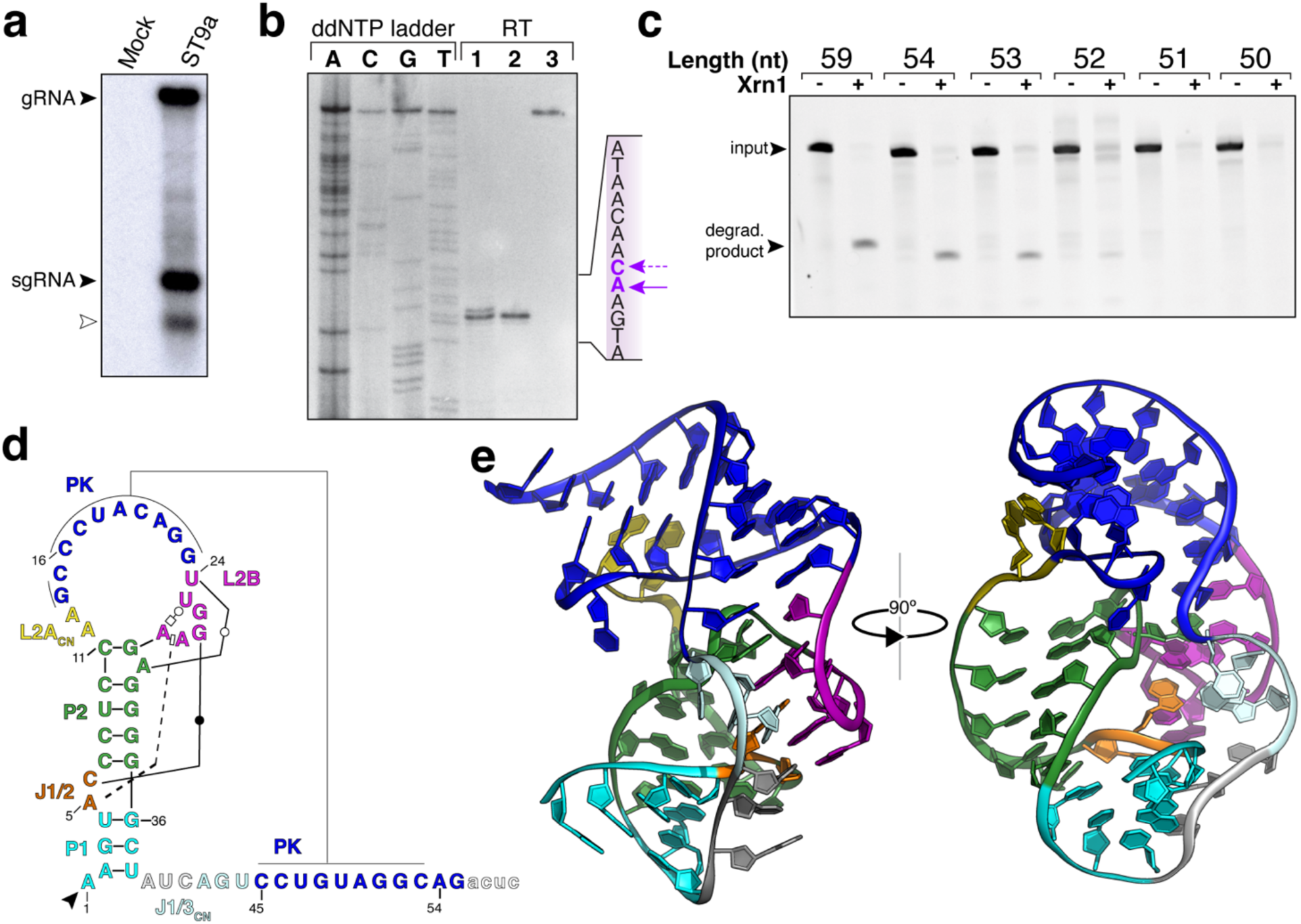
A compact RNA structure in the 3′ UTR of the tombusvirus-associated RNA ST9a blocks degradation by Xrn1. a) Northern blot of total RNA from *N. benthamiana* mock-infected or infected with ST9a for 8 days. Probes are against the ST9a 3′ UTR. The white arrow denotes an unspecified RNA species, likely due to additional in-cell processing of the sgRNA or host rRNA trapping. b) Sequencing gel of cDNA from Xrn1 degradation reaction (1), crystallization construct (2) and undigested RNA (3) to map the Xrn1 stop site at single nucleotide resolution. Dideoxy sequencing lanes are labeled and used to determine the sequence around the Xrn1 stop site, depicted on the right with arrows. c) *In vitro* Xrn1 degradation reaction of ST9a 3′ UTR to determine the minimal Xrn1-resistant element. Reactions were resolved by dPAGE and visualized by ethidium bromide staining. Length in nucleotides (nt) corresponds to the resistant product, counting from the Xrn1 stop site as determined in b. d) Secondary structure diagram of the ST9a xrRNA crystallization construct. Non– Watson–Crick base pairs are in Leontis–Westhof annotation^63^ and the Xrn1 stop site is depicted by the arrow. Non-modeled nucleotides are shown as lowercase letters. e) Ribbon diagram of the ST9a xrRNA crystal structure. Colors match d.

### The ST9a xrRNA adopts a class 3 fold with an extended pseudoknot

To dissect the molecular architecture of the ST9a xrRNA, we solved the structure of the minimal resistant element to 2.9 Å resolution using x-ray crystallography (Fig. 1d-e, Extended Data Fig. 3d-e, Extended Data Fig. 4, see Materials and Methods for details). The asymmetric unit of the crystal contained two almost identical RNA monomers with a root-mean-square deviation (RMSD) of 0.237 Å (Extended Data Fig. 4b-c). As predicted, the RNA adopts a class 3 xrRNA conformation with a SL of coaxially stacked helices P1 and P2 (Fig. 1e, Extended Data Fig. 3d-e). For the first time, a PK is captured *in cis* between the apical loop and the 3′ end of each monomer, leading to the characteristic protective ring encircling the 5′ end of the RNA element (Fig. 2a, Extended Data Fig. 5a-c). A 9 base pair PK is formed by a stretch of eight consecutive Watson-Crick base pairs between nucleotide C16 to G23 in the apical loop and C45 to G52 in the 3′ end, extended by one additional base pair between G14 and C53 next to flipped-out C15 (Fig. 2a-b). The PK forms a continuous stack with L2B, a highly conserved hairpin structure embedded in the apical loop, creating an extended stacking network that makes one full helical turn (Fig. 2b). Mutations to the central nucleotides of the PK (U47A/G48C/U49A) disrupt Xrn1-resistance *in vitro* and sgRNA formation during ST9a infection of *N. benthamiana* (Fig. 2c-d, Extended Data Fig. 2). These defects can be rescued by compensatory mutations to the apical loop that restore the PK (Fig. 2c-d, Extended Data Fig. 2), demonstrating that, as in other class 3 xrRNA variants^40–42^, PK formation is essential for exoribonuclease resistance. Mutating the PK during ST9a infection also significantly reduces genomic RNA (gRNA) accumulation, suggesting that xrRNAs play an important role in ST9a replication (Fig. 2d, Extended Data Fig. 2). Surprisingly, shortening of the PK eliminates Xrn1-resistance *in vitro* (Fig. 2c). This contrasts other class 3 xrRNA variants, where a 3 to 5 base pair PK is sufficient to block Xrn1^40,41^. The strong reliance of the ST9a xrRNA on an extended PK may have contributed to our ability to crystallize the RNA in its closed conformation. The PK is further stabilized by minor groove interactions of A12 and A13 in the apical loop (Fig. 2e-f). These A-minor interactions are essential, as mutating both adenosines to uracil (A12U/A13U) or deleting one adenosine (delA13) disrupts Xrn1 resistance (Fig. 2e). However, A12 and A13 are not conserved in other class 3 xrRNAs; instead, they are part of the highly variable L2A sequence in the apical loop^41,42^. We propose that L2A variability has evolved to accommodate length variations within P2 and the PK, helping to maintain the overall compact xrRNA structure. A similar strategy was observed in the PLRV xrRNA, where flipped-out nucleotides in the L2A region allow for the addition of extra nucleotides compared to the SCNMV xrRNA (Extended Data Fig. 5d)^42^.

**Figure 2.**
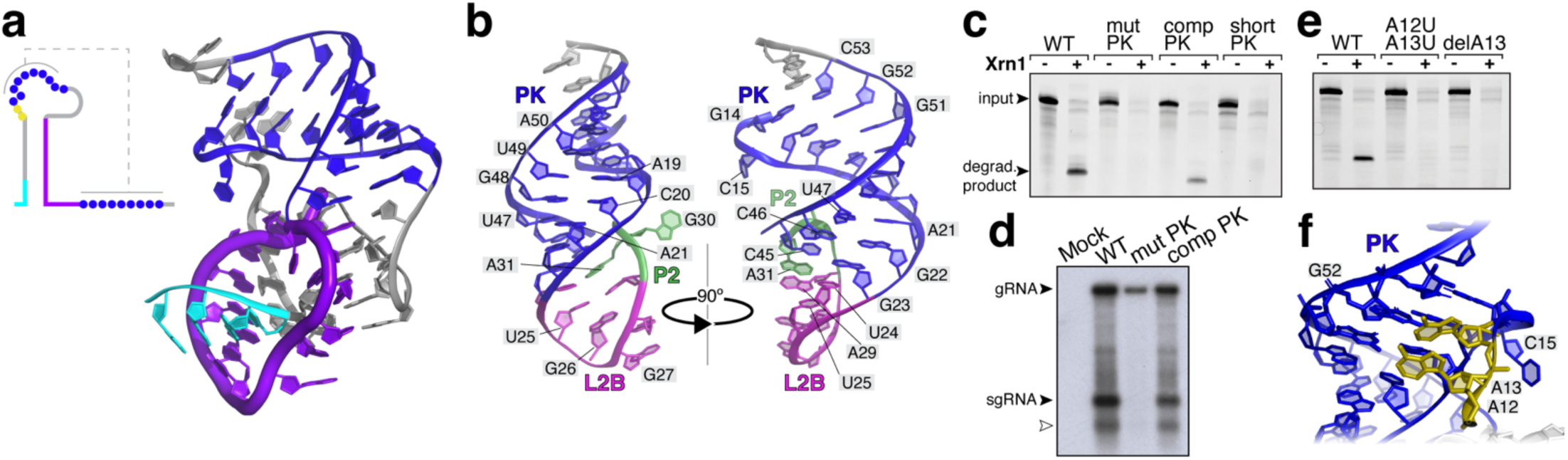
The structure of the ST9a xrRNA contains a 9 base pair PK and a protective ring around the 5′ end. a) Ribbon diagram (right) and 2D structure diagram (left) of the ST9a xrRNA structure with the 9 base pair PK in blue, the 5′ end in cyan and the protective ring around the 5′ end in purple. b) Coaxial stacking of the PK and L2B region. Colors match Fig. 1d-e. c) *In vitro* Xrn1 degradation reaction of ST9a xrRNA wild type (WT) RNA and the indicated mutants. Reactions were resolved by dPAGE and visualized by ethidium bromide staining. d) Northern blot of total RNA from *N. benthamiana* mock-infected or infected with ST9a WT or the indicated mutants. Probes are against the ST9a 3′ UTR. The white arrow denotes an unspecified RNA species, likely due to additional in-cell processing of the sgRNA or host rRNA trapping. e) *In vitro* Xrn1 degradation reaction of ST9a xrRNA WT RNA and the indicated mutants. Reactions were resolved by dPAGE and visualized by ethidium bromide staining. f) Details of the interactions between A12 and A13 with the minor groove of the PK.

### A conserved long-range interaction network stabilizes the characteristic ring topology of the xrRNA

A PK is necessary but not sufficient for xrRNA’s resistance to exoribonuclease degradation. The ST9a xrRNA structure reveals additional conserved tertiary interactions that are crucial for maintaining the protective ring topology. Notably, we identify a core network of long-range interactions involving four conserved motifs centered around the apical loop protrusion L2B (Fig. 3a-b). This network buttresses the 3′ face of the protective ring, likely contributing to the RNA’s stability against the pulling forces of exoribonucleases. The L2B backbone makes a sharp turn, resembling a U-turn RNA tetraloop^49^, but it is closed by a Hoogsteen base pair between U25 and A29 and a reverse Watson-Crick base pair between U24 and A31, with A31 extruding from P2 (Fig. 3a-b). This motif serves as a docking platform for A5 and C6 in the J1/2 bulge (Fig. 3c). A5 stacks between A28 and A29 and forms two hydrogen bonds with the sugar edge of G26, while C6 forms a Watson-Crick base pair with G27 (Fig. 3c). Interestingly, the C6-G27 interaction replaces a conserved G-C base pair seen in most other class 3 xrRNAs^40,41^, making it the first observed instance of covariation at this position. While this base pair stabilizes the J1/2-L2B tertiary interaction, its importance varies across xrRNA variants. For example, G8-C27 pairing is essential for exoribonuclease resistance in PLRV^42^, but mutating the same interaction in ST9a has only a minor effect on exoribonuclease resistance (Extended Data Fig. 5e-f), and SCNMV lacks the base pair altogether^40^. In contrast, mutations to any other nucleotide of the core network completely abolish exoribonuclease resistance *in vitro*, and an A5U mutation in ST9a prevents sgRNA production during infection (Fig. 3d-e, Extended Data Fig. 5g-h). As with PK mutations (Fig. 2d), the A5U mutation reduces gRNA accumulation in infected cells (Fig. 3d).

**Figure 3.**
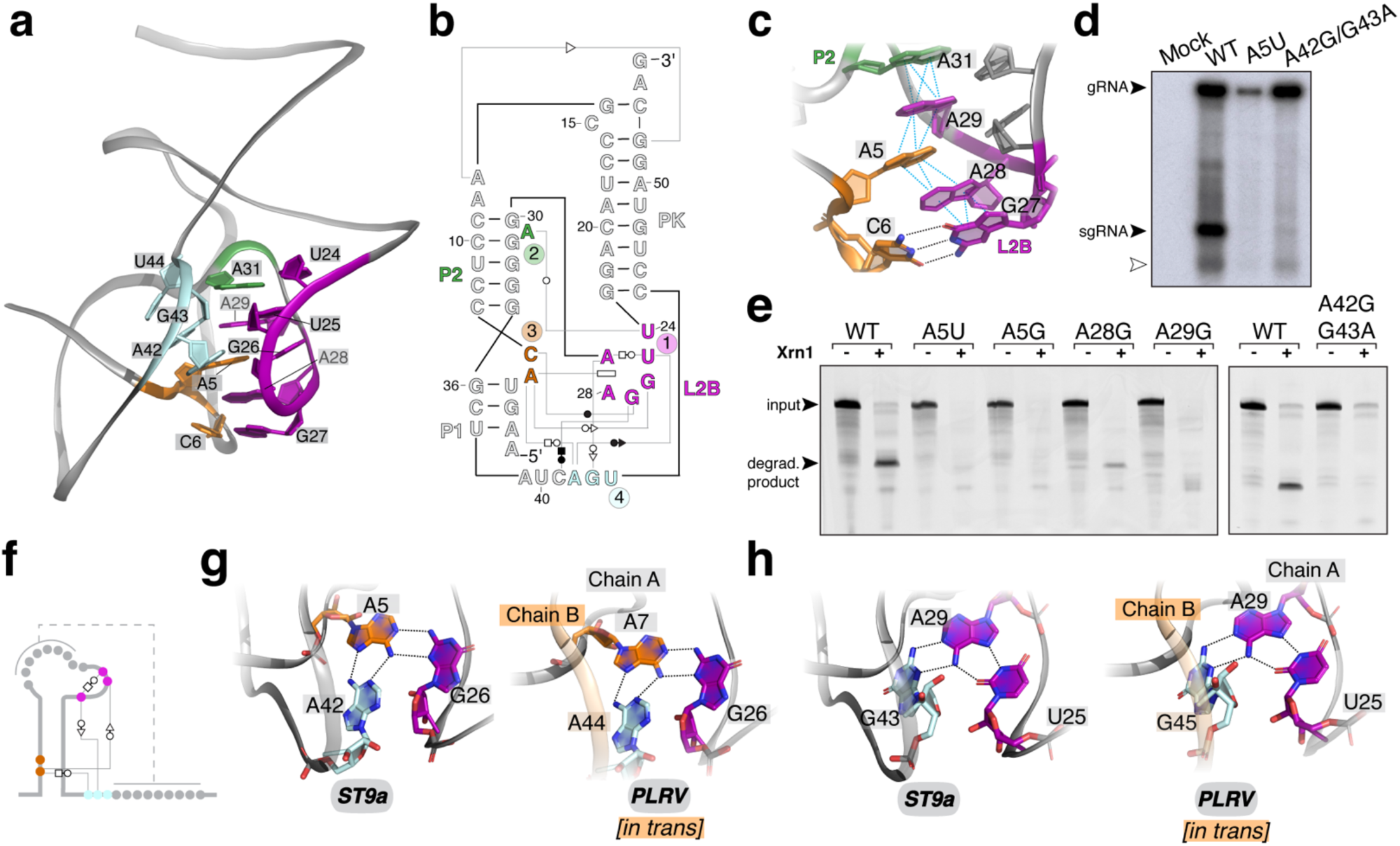
A conserved interaction network forms the core of the xrRNA structure. a) Details of the tertiary interaction network centered around the L2B region with colors to match Fig. 1d-e. b) 2D diagram of the tertiary interaction network depicted in a. Non–Watson–Crick base pairs are in Leontis–Westhof annotation^63^. c) Details of the L2A pi-stacking network with colors to match a. d) Northern blot of total RNA from *N. benthamiana* mock-infected or infected with ST9a WT or the indicated mutants. Probes are against the ST9a 3′ UTR. The white arrow denotes an unspecified RNA species, likely due to additional in-cell processing of the sgRNA or host rRNA trapping. e) *In vitro* Xrn1 degradation reaction of ST9a xrRNA WT RNA and the indicated mutants. Reactions were resolved by dPAGE and visualized by ethidium bromide staining. f-h) Conserved base triple interactions, schematically depicted in (f), centered around A5 (g) and A29 (h) are part of the conserved ST9a xrRNA core. Identical hydrogen bonds are formed *in cis* in ST9a xrRNA (left) and *in trans* between 2 RNA molecules in PLRV xrRNA (right, PDB: 7JJU^42^).

The L2B-centered core network is highly conserved across class 3 xrRNAs and adopts almost identical conformations in all crystal structures (RMSD of 0.337 Å between the ST9a and PLRV L2B and 0.596 Å between the ST9a and SCNMV L2B) (Extended Data Fig. 5c)^40,42^. This similarity is remarkable considering that PLRV and SCNMV xrRNAs were crystallized as dimers without the PK formed *in cis*^40,42^ (Extended Data Fig. 5a-b). In these structures, the same J1/2-L2B long range interactions that position the protective ring in ST9a hold the SL in a tilted conformation, poised to transition into the PK state^40,42^. This poised SL is likely integral to the rapid co-degradational remodeling of the xrRNA^40^. The only nucleotides of the long-range xrRNA core network not observed *in cis* in previous crystal structures are A42 and G43. These nucleotides, part of a highly conserved AGY motif^41,42^, form the 3′ end of the protective ring (Fig. 3a-b, 3f). Through non-canonical base triple interactions with A5 and L2B, they anchor the ring to the core of the xrRNA. A42 interacts with the Hoogsteen face of A5 to form an A42-A5-G26 base triple (Fig. 3g). Additionally, the sugar edge of G43 forms base triple interactions with A29 of the U25-A29 Hoogsteen base pair (Fig. 3h). While these interactions were not observed *in cis* in earlier structures, identical base triple interactions were established as part of the crystallographic dimers of both PLRV and SCNMV xrRNA^40,42^ (Fig. 3g-h), further supporting the notion that the L2B network forms the conserved core of class 3 xrRNA structures. As with all other nucleotides in the core, mutating A42 and G43 completely abolishes exoribonuclease resistance *in vitro* and severely reduces sgRNA production during infection (Fig. 3d-e).

### Magnesium ions stabilize the xrRNA structure

Magnesium ions (Mg^2+^) play crucial roles in stabilizing tertiary RNA structures, and the mechanical anisotropy of flavivirus xrRNAs is highly Mg^2+^-dependent^24,50^. To assess ST9a xrRNA’s dependence on Mg^2+^, we performed quantitative *in vitro* RNA degradation assays across a range of Mg^2+^ concentrations. Robust Xrn1-resistance was observed at concentrations of ≥ 2.5 mM Mg^2+^, with resistance dropping below 30% at 1 mM Mg^2+^ (Fig 4a, Extended Data Fig. 6a). This suggests that ST9a xrRNA forms a stable, exoribonuclease-resistant fold at near-physiological Mg^2+^ concentrations. Consistent with a Mg^2+^-driven stabilization of the xrRNA tertiary fold, we observed Mg^2+^-dependent changes in thermal stability of ST9a xrRNA in UV melting experiments: Two melting events observed in the absence of Mg^2+^ merged into a single high-temperature melting event (79.16°C) in the presence of Mg^2+^ (Fig. 4b).

**Figure 4.**
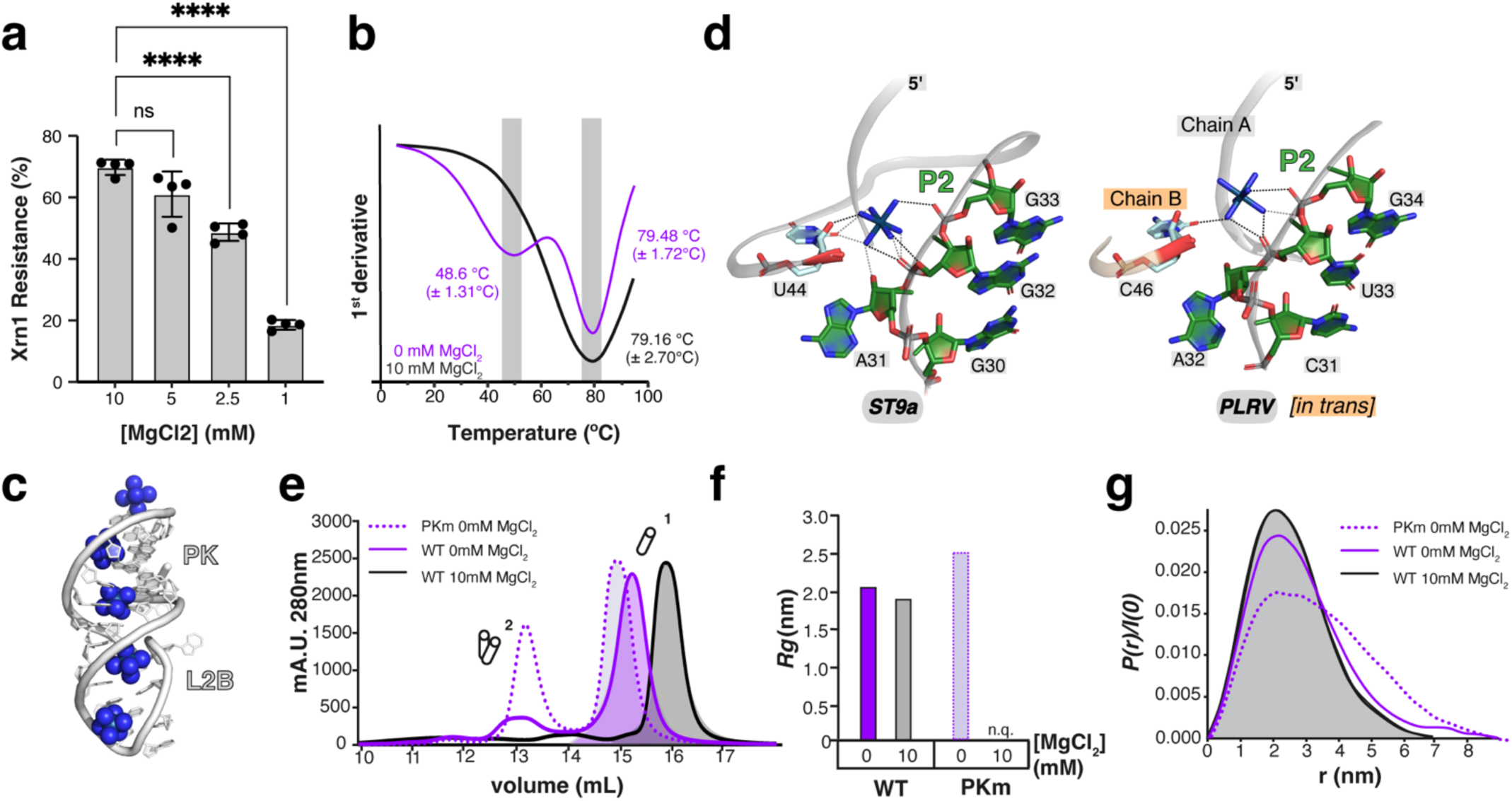
Coordinated metal ions stabilize the xrRNA structure. a) Quantitative Xrn1 degradation assay with 3′-^32^P-labeled ST9a xrRNA at varying Mg^2+^ concentrations. n=4. Error bars show SD. **** adjusted P- value <0.0001. b) Thermal melting curves of the ST9a xrRNA at the indicated Mg^2+^ concentrations. Shown is the first derivative of melting curves at 266 nm. Samples were measured in biological replicates (n=3). c) Iridium(III) hexammine ions in the major groove of the L2B-PK helix. d) Details of a conserved iridium(III) hexammine binding site in the ST9a xrRNA structure (left) and PLRV xrRNA structure (right). Note that the iridium(III) hexammine ion is coordinated by nucleotides from two RNA molecules of the crystallographic dimer in the PLRV xrRNA. e) Zoom-in to SEC profiles of the indicated ST9a xrRNA constructs at 0 or 10 mM Mg^2+^. f) Plot of *Rg* and g) scattering-pair distance distribution (*P*(*r*)) profiles derived from SAXS data for the monomeric peaks from e, showing Mg^2+^ induced compaction. See also Extended Data Figures 6 and 7 and Table 2.

**Table 1.** Data collection and refinement statistics.

|  |  |
| --- | --- |
| **Structure** | ST9a xrRNA |
| **Beamline** | 19-ID (NYX) NSLS-II |
| **Wavelength (Å)** | 0.9795 |
| **Resolution range** | 41.06 - 2.9 (3.004 - 2.9) |
| **Space group** | C 1 2 1 |
| **Unit cell** | 107.637 42.392 84.491 90 103.618 90 |
| **Total reflections** | 31177 (3223) |
| **Unique reflections** | 8421 (494) |
| **Multiplicity** | 3.7 (3.8) |
| **Completeness (%)** | 87.25 (58.81) |
| **Mean I/sigma(I)** | 10.53 (1.71) |
| **Wilson B-factor** | 55.77 |
| **R-merge** | 0.101 (0.8365) |
| **R-meas** | 0.1177 (0.9738) |
| **R-pim** | 0.05966 (0.4913) |
| **CC1/2** | 0.981 (0.817) |
| **CC\*** | 0.995 (0.948) |
| **Reflections used in refinement** | 7384 (494) |
| **Reflections used for R-free** | 739 (50) |
| **R-work** | 0.2148 (0.2591) |
| **R-free** | 0.2575 (0.3468) |
| **CC(work)** | 0.818 (0.759) |
| **CC(free)** | 0.737 (0.706) |
| **Number of non-hydrogen atoms** | 2503 |
| **macromolecules** | 2333 |
| **ligands** | 170 |
| **solvent** | 0 |
| **Protein residues** | N/A |
| **RMS(bonds)** | 0.002 |
| **RMS(angles)** | 0.50 |
| **Ramachandran favored (%)** | N/A |
| **Ramachandran allowed (%)** | N/A |
| **Ramachandran outliers (%)** | N/A |
| **Rotamer outliers (%)** | 0.00 |
| **Clashscore** | 8.41 |
| **Average B-factor** | 52.60 |
| **ligands** | 72.18 |
| **Number of TLS groups** | 8 |
| **PDB code** | 9CFN |

2 Statistics for the highest-resolution shell are shown in parentheses.

**Table 2.**
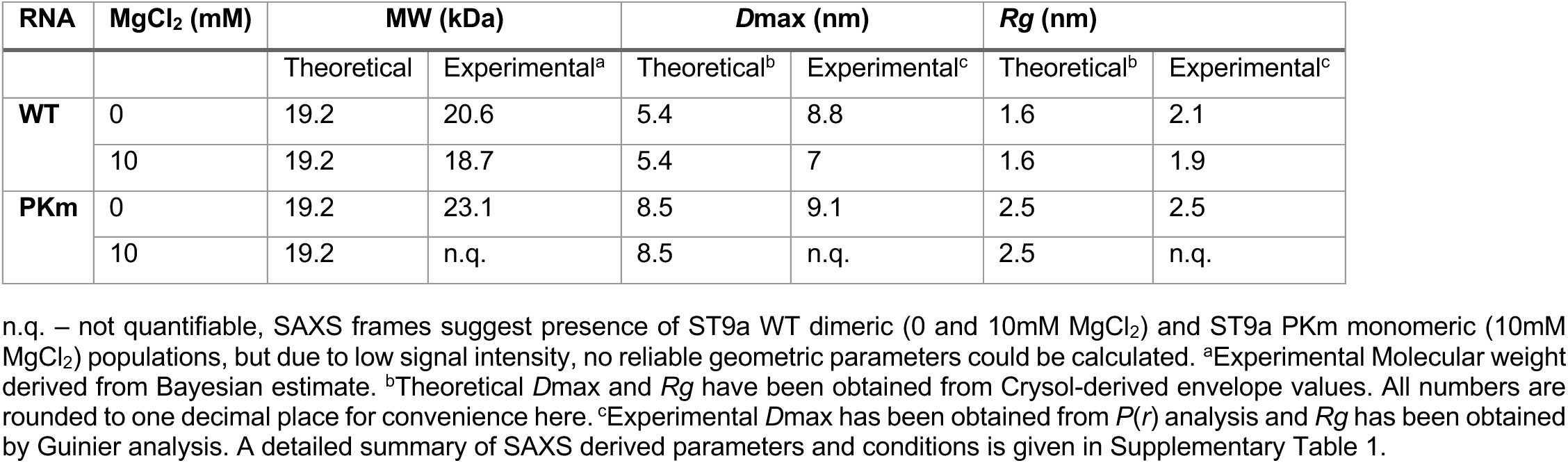
Geometric parameters derived from SEC-SAXS experiments.

ST9a xrRNA crystals were grown in a buffer containing MgCl_2_ and later soaked in iridium(III) hexammine chloride for experimental phasing by single-wavelength anomalous diffraction^51^ (see Methods). This approach allowed us to investigate metal ion binding sites at high resolution. We identified one Mg^2+^ per RNA molecule, coordinated by U40, C41, and the conserved G43, suggesting a role in positioning the protective ring around the 5′ end (Extended Data Fig. 6b-c). Since iridium(III) hexammine is isosteric to hexahydrated Mg^2+^, its positions in the crystal structure serve as proxies for Mg^2+^ binding sites^51^. A total of 24 iridium(III) hexammine ions were identified in the asymmetric unit (Extended Data Fig. 4e). Notably, 14 of these are located in the major grooves of coaxially stacked helices L2B-PK and P1-P2, resembling the position of hexahydrated Mg^2+^ in A-form RNA^52^ (Fig. 4c). Several iridium(III) hexammine ions cluster near the 5′ end of the protective ring, likely neutralizing the charges from closely positioned backbone phosphates in the tightly wrapped structure (Extended Data Fig. 4e). Most strikingly, one iridium(III) hexammine ion is positioned between the 5′ and 3′ ends of the protective ring, coordinated by the backbone phosphates of G32 and G33, along with a hydrogen bond network involving the ribose and base of U44, such that the ion is effectively sealing the ring around the 5′ end of the RNA structure (Fig. 4d). Similar interactions are formed in the crystal structures of PLRV and SCNMV xrRNAs, where they connect two RNA molecules *in trans* (Fig. 4d)^40,42^. The conservation of these interactions supports the idea that metal ion-dependent sealing of the ring is a critical feature of the class 3 xrRNA topology.

### The ST9a xrRNA adopts a compact structure in solution

To assess whether the compact monomeric xrRNA structure observed by crystallography is preserved in solution, we performed small-angle X-ray scattering (SAXS) coupled with size-exclusion chromatography (SEC)^53^ to evaluate the global shape and organization of monodispersed xrRNA populations (Fig. 4e-g, Table 2, Extended Data Fig. 6d-g, Extended Data Fig. 7a-g, Supplementary Table 1). Even at high RNA concentrations used for SEC-SAXS (3 mg/ml), wild type ST9a xrRNA remained predominantly monomeric in solution as indicated by the SEC elution profile and SAXS-derived molecular weights (Fig. 4e, Table 2). A minor dimer population was observed in the absence of Mg^2+^, but the addition of Mg^2+^ stabilized the monomeric form and induced compaction of the monomer as reflected by a delayed SEC elution time (Fig. 4e, Extended Data Fig. 6d-e) and a reduction in the radius of gyration (*Rg*) from 2.1 to 1.9 nm (Fig. 4f, Table 2). The scattering pair distance distribution (*P*(*r*)) profiles further demonstrate that the ST9a xrRNA monomer adopts a globular fold, which is significantly compacted in the presence of Mg^2+^, leading to a 2 nm reduction in maximum dimension (*Dmax*) (Fig. 4g, Table 2). The experimental SAXS data for the monomeric ST9a xrRNA are in good agreement with calculated curves based on the crystal structure (Extended Data Fig. 7a-d), with the best fit observed when SAXS data for a 54-nucleotide minimal xrRNA element were collected in Mg²⁺-containing buffer (χ² = 1.26) (Extended Data Fig. 7c). This suggests that the crystal structure accurately represents a major population in solution. To further validate these findings, we analyzed an ST9a xrRNA variant with a mutated PK (PKm). This mutation led to an increased dimer population (Fig. 4e, Extended Data Fig. 6f), which was further amplified by Mg^2+^ (Extended Data Fig. 6g). Additionally, the monomeric fraction of ST9a-PKm exhibited an expanded geometry in solution as indicated by an increase in *Rg* (Fig. 4f), a broader spread in the *P*(*r*) curve (Fig. 4g), and reduced structural compactness seen in the dimensionless Kratky plot (Extended Data Fig. 7g). Calculated SAXS curves for ST9a xrRNA PKm modeled in an open conformation using RNAmasonry^54^ best fit the experimental SAXS data of ST9a-PKm (χ² = 0.96) (Extended Data Fig. 6f).

### Class 3 xrRNA variants are widespread in plant infecting viruses

Previous computational searches for xrRNA structures with conserved sequence and structural features identified over 50 distinct members of the xrRNA class 3 but notably overlooked the ST9a xrRNA^41,42^. This omission is likely due to the high stringency of the search algorithm, combined with unique features of the ST9a xrRNA, such as its longer PK and the C6-G27 covariation of a typically conserved G-C base pair (Fig. 3c). By incorporating the new sequence and structural insights from this study, we refined the search model and identified 363 high-confidence hits representing unique class 3 xrRNA variants (Fig. 5a, Extended Data Fig. 8, Supplementary Table 2). These newly identified variants span several families of plant-infecting +RNA viruses, with the majority found in the *Solemoviridae* family (Fig. 5b). Many of the xrRNA-containing viruses were previously annotated as *Luteoviridae*^41^, but have recently been re-classified as *Solemoviridae* and *Tombusviridae*^55^. Interestingly, most of these new putative xrRNAs are in the intergenic regions of viral genomes (Fig. 5c), suggesting a role in the production of protein-coding sgRNAs, potentially facilitated by the cap-independent translation mechanisms commonly used by plant-infecting RNA viruses^56^. Notably, we also identified the first putative class 3 xrRNAs in the 5′ UTR and coding sequences (CDS) (Fig. 5c), although their functional relevance remains to be confirmed. Overall, the identification of hundreds of related xrRNA structures across diverse viral genomes – with potential roles in producing both protein-coding and non-coding sgRNAs – underscores the importance of this expanding class of functional structured RNAs.

**Figure 5.**
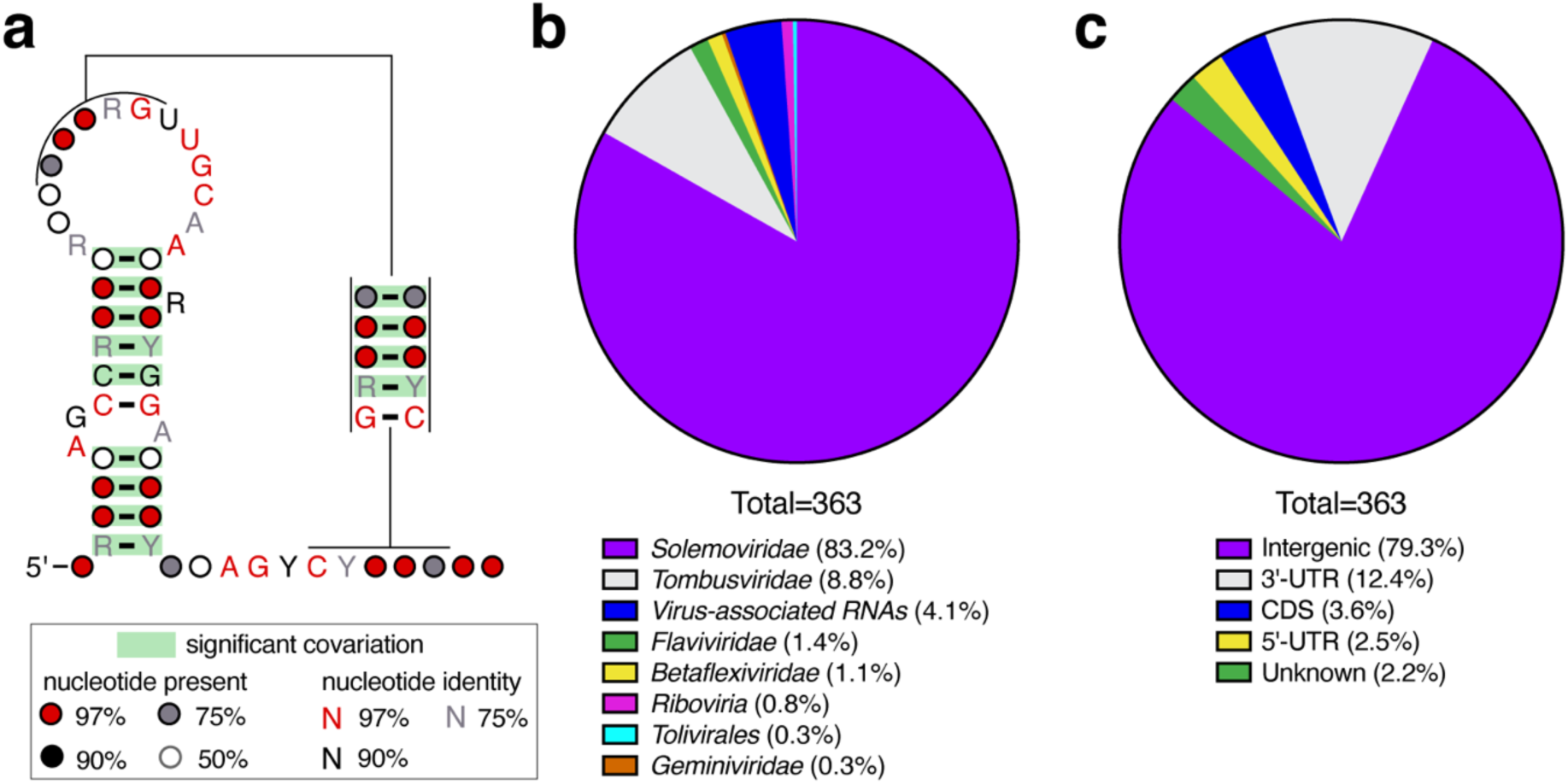
Widespread distribution of class 3 xrRNAs in plant-infecting RNA viruses. a) Consensus sequence secondary structure of class 3 xrRNAs based on a comparative sequence alignment of 363 sequences of plant-infecting RNA viruses and virus-associated RNAs. Y = pyrimidine; R = purine. b) Distribution of xrRNA sequences from (a) across viral genera. Note that many xrRNA-containing viruses previously annotated as *Luteoviridae*, have been re-classified as *Solemoviridae*^55^. c) Metagenome analysis of xrRNA sequences from (a).

### The ST9a xrRNA structure reveals a common molecular mechanism to block exoribonucleases

A defining feature of all viral xrRNA structures is their characteristic slipknot-like topology with a central protective ring through which the 5′ end passes^12^. Our high-resolution structure of a class 3 xrRNA in its active conformation shows that its ring closely resembles that of the well-characterized class 1 xrRNAs, such as those from Murray Valley encephalitis (MVE)^11^ virus and ZIKV^22^. To compare these two highly distinct xrRNA classes, we aligned the central ring of ST9a xrRNA to ZIKV xrRNA1^22^, which served as a model of the class 1 xrRNAs (Fig. 6a-c). In both cases, ring formation relies on direct interactions between the 5′ end and ring nucleotides, ensuring the ring closes only when the 5′ end is positioned at its center (Fig. 6d). In the ZIKV xrRNA, this positioning is achieved through a base triple and an interwoven pseudoknot (PK1) formed between the 5′ end and the 3WJ (Fig. 6d)^11,22^. In contrast, the ST9a xrRNA accomplishes the same outcome using its P1 helix (Fig. 6d). The Xrn1 stop site at the base of P1 in the ST9a xrRNA (Fig. 1b, 1d) suggests that P1 is partially unwound when the 5′ end enters Xrn1’s narrow RNA entry channel, meaning P1 does not directly contribute to the RNA’s mechanical stability. Rather, stability is provided by the protective ring, which is constructed from several helices and stabilized by long-range interactions that buttress the 3′ face of the ring to redistribute the enzyme’s pulling forces. In ZIKV, the ring is formed by helices P1 and P3^22^, while in ST9a the ring is created by helix P2 and the long-range L2B interaction network (Fig. 6e). Notably, P2 in ST9a and P3 in ZIKV xrRNA occupy nearly identical positions, anchoring the 5′ side of the ring in both structures (Fig. 6e). However, the 3′ side of the ring is stabilized differently: ZIKV xrRNA uses P1 stacking on P2, while ST9a xrRNA relies on its network of long-range interactions centered around L2B (Fig. 3a-b, 6f). In both classes, the ring is sealed by a reverse Watson-Crick base pair (A37-U51 in ZIKV xrRNA and A31-U24 in ST9a xrRNA), creating a concave ring to form an effective brace against an exoribonuclease approaching from the 5′ end (Fig. 6a, c). The different strategies used by class 1 and class 3 xrRNAs to stabilize the remarkably similar ring highlight how viruses have evolved distinct structural solutions to blocking an RNA degrading enzyme (Supplementary video 1).

**Figure 6.**
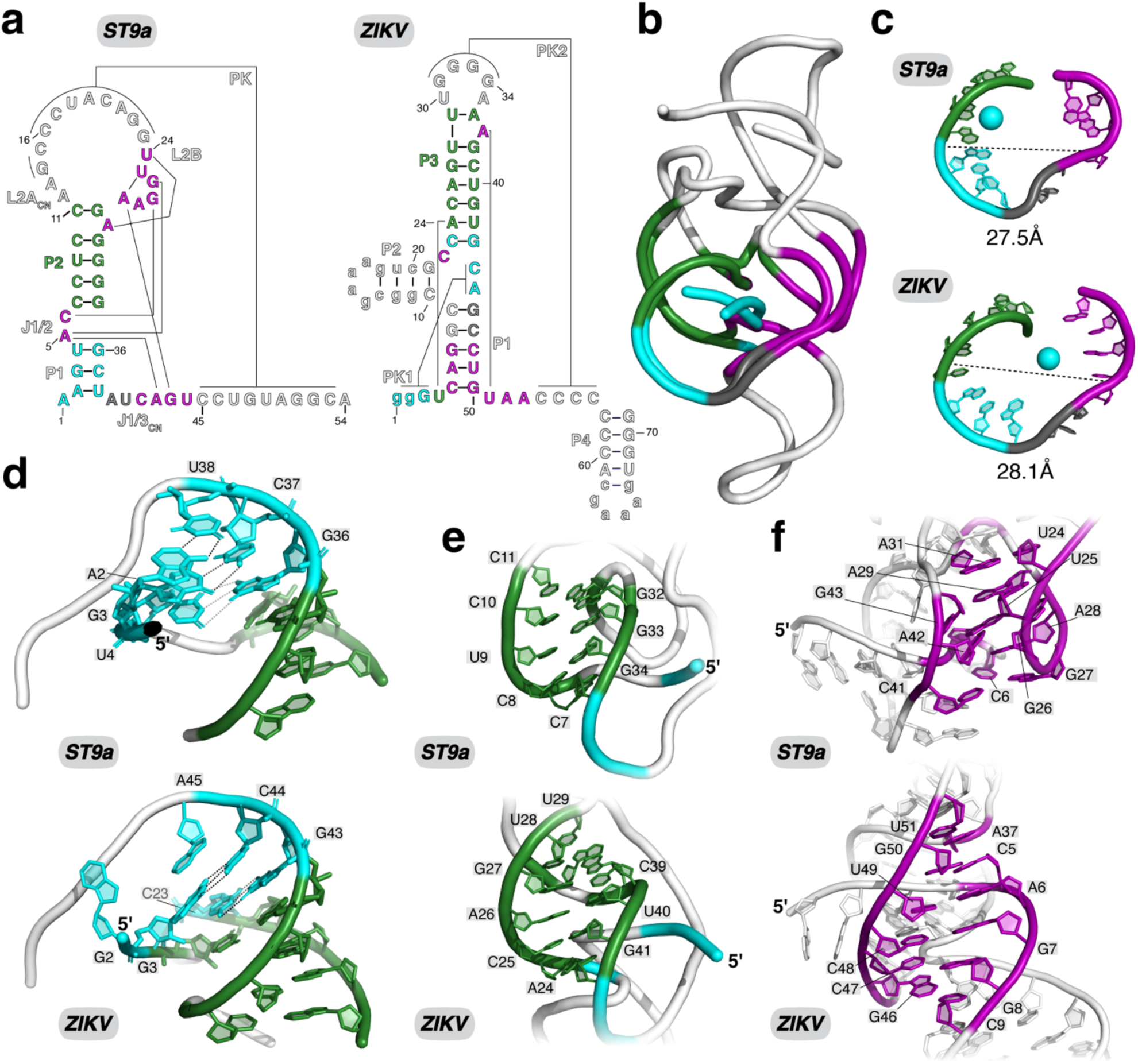
Structural comparison of class 1 and class 3 xrRNAs uncovers a conserved molecular mechanism to block host exoribonucleases. a) 2D structure of the ST9a (left) and ZIKV (right) xrRNAs. Colors indicate common features. Lowercase letters in ZIKV xrRNA denote sequences that were mutated to aid crystallization. b) Overlay of the core of ST9a and ZIKV xrRNA with colors to match a. c) The protective rings of ST9a (top) and ZIKV (bottom) xrRNAs have similar dimensions. Colors to match a. The cyan sphere indicates the position of 5′ end. d) Details of the nucleotides positioning the 5′ end within the center of the ring of ST9a (top) and ZIKV (bottom) xrRNA. e) Details of the helix buttressing the 5′ side of the protective ring in ST9a (top) and ZIKV (bottom) xrRNA. f) Details of the unique interaction networks stabilizing the 3′ side of the protective ring in ST9a (top) and ZIKV (bottom) xrRNA.

## Discussion

The high-resolution structure of a class 3 xrRNA in its closed, active conformation marks a significant advance in our molecular understanding of programmed exoribonuclease-resistance. Strikingly, the ST9a PK conformation exhibits minimal structural differences to previously solved structures of a class 3 xrRNA in the open SL conformation^40,42^, with only a few interactions needing rearrangement during PK formation (Extended Data Fig. 5a-c). This observation suggests that the open conformation is already pre-configured for the conformational switch to the active PK state, laying the groundwork for a hierarchical folding model: P1 initially positions the 5′ end (Fig. 6d), followed by anchoring of the AGY motif to the L2B network (Fig. 3g-h, Fig. 6f), magnesium-dependent sealing of the ring (Fig. 4d), and finally stabilization by PK formation (Fig. 2a-b). While P1 plays a critical role in directing this folding path (Fig. 6d) and the ST9a xrRNA structure shows that P1 and PK can coexist in the same conformation (Fig. 1e), the Xrn1 stop site at the 5′ base of P1 implies partial P1 unwinding upon degradation. This suggests that P1 does not directly contribute to the mechanical stability of the fully formed xrRNA. Notably, earlier studies on class 3 xrRNAs with shorter PKs revealed highly dynamic behavior, with these RNAs frequently transitioning between SL and PK conformations^40^. This observation led to the hypothesis that co-degradational refolding–where the PK forms during Xrn1 degradation–is the basis for robust resistance to exoribonuclease activity^40^. Whether ST9a xrRNA and related xrRNAs with significantly longer PKs display similar dynamics or adopt a more stable closed conformation–as was proposed for flaviviral class 1 xrRNAs^11,12,24,26,50^ – remains an open question. It is equally unknown how this conformational ensemble might be regulated inside cells^57,58^. Interestingly, flaviviral class 1 xrRNAs appear to rely on a similar hierarchical folding pathway to position the 5′ end within the protective ring^12^. However, in class 1 xrRNAs, this positioning is facilitated by PK1 (Fig. 6d)^11,22^, highlighting the distinct strategies used by different xrRNA classes to solve a common folding challenge: how to thread the 5′ end through a closed ring within the viral genome context (Supplementary video 1).

Structural comparisons of class 1 and class 3 xrRNAs reveal that the characteristic ring can be embedded in highly distinct scaffolds. This variability underscores the remarkable versatility of xrRNAs in achieving a common functional outcome through structurally diverse mechanisms (Fig. 6). While flaviviral xrRNAs have been categorized into class 1a, class 1b and class 2 based on variations in 2D and 3D structures^13^ (Extended Data Fig. 1b), all flaviviral xrRNAs are related and share key features, such as the 3WJ that organizes the ring topology^11,13,17,22,23,59^–flavivirus xrRNA classes can thus be considered variations of a single structural mechanism. In contrast, class 3 xrRNAs employ a fundamentally different strategy to form a similar protective ring topology. Indeed, class 3 xrRNAs differ so significantly from other xrRNA classes that it remains unknown whether they evolved from a common ancestor or through convergent evolution. Given the vast number of viral RNA sequences with unknown functions, it is highly likely that additional mechanisms for forming and stabilizing the conserved ring exist, motivating the search for novel xrRNA classes. Recent discoveries of putative xrRNAs in the genomes of viruses from the *Arenaviridae, Benyviridae, Bunyaviridae, Betaflexiviridae, Tombusviridae* and *Virgaviridae* families^60–62^ support this notion. Based on predicted secondary structures, none of these putative new xrRNAs belong to the three previously characterized xrRNA classes, suggesting that they likely employ entirely new strategies to create exoribonuclease-resistant structures. Exploring xrRNA variability promises to expand our mechanistic understanding of programmed exoribonuclease resistance and to uncover broadly generalizable features of xrRNAs. Structure-guided searches could also reveal xrRNAs in new contexts, potentially expanding the role of programmed exoribonuclease resistance beyond the realm of +RNA viruses.

Finally, while significant progress has been made in understanding the structure and dynamics of these viral RNA elements, much less is known about their biological functions. The finding that most class 3 xrRNAs are in intergenic regions of viral genomes suggests they may serve roles distinct from those of flaviviral class 1 and class 2 xrRNAs, which have so far been exclusively identified in the 3′ UTRs. High-resolution structural insights into the class 3 xrRNA fold enabled the precise design of point mutations that disrupt exoribonuclease resistance. Our initial findings demonstrate that ST9a with a compromised xrRNA is attenuated during infection (Fig. 2d, Fig. 3d). Future studies using similar mutants will be essential to systematically investigate the functional roles of these elements during the viral infection cycle.

## Acknowledgements

We thank Kevin Battaile for assistance at the NYX beamline and Shirish Chodankar for assistance at the LiX beamline at NSLS-II. We also thank Prof. Andreas Schlundt and Karthikeyan Dhamotharan for valuable discussion regarding SAXS data, and Steve Bonilla and members of the Steckelberg lab for carefully reading the manuscript and providing valuable feedback.

