## Supplementary Material for "The pseudoknot structure of a viral RNA reveals a conserved mechanism for programmed exoribonuclease resistance"

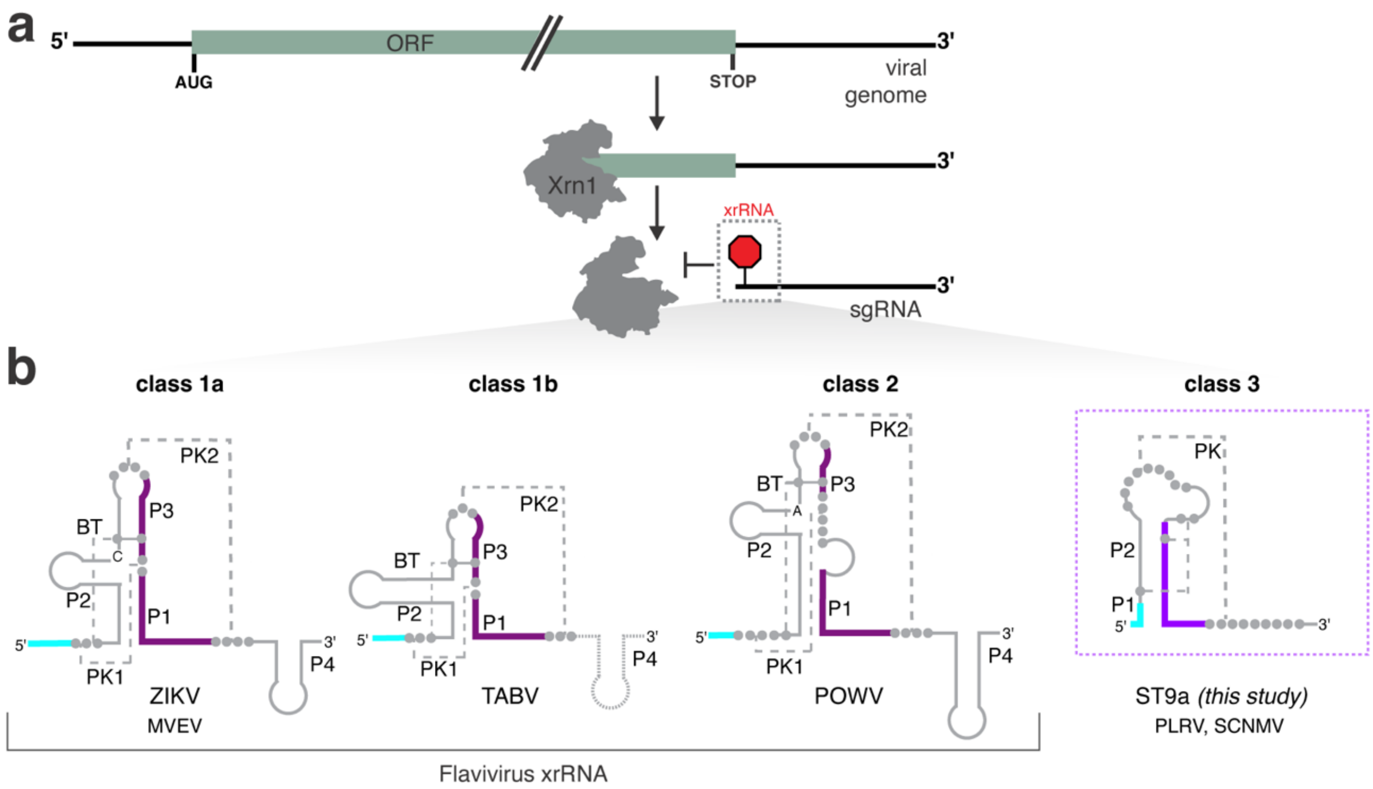


**Extended Data Fig. 1. Diversity of viral xrRNAs.** a) Schematic representation of viral programmed exoribonuclease resistance. Cellular exoribonucleases, such as Xrn1, degrading the viral genome with 5′-3′ directionality are stopped by exoribonuclease-resistant RNA structures called xrRNAs. This trims the viral genome to produce sgRNAs with important functions during viral infection[1]. b) Schematic representation of different xrRNA classes identified to date. The flaviviral xrRNAs (left) all rely on two PKs centered around a 3WJ. Based on sequence and structure variation, flaviviral xrRNAs can be subdivided into class 1, class 1b and class 2 xrRNAs. Class 3 xrRNAs (right) are found in plant-infecting RNA viruses and use a single PK to form an exoribonuclease-resistant structure. Names under the 2D diagrams list members of the respective classes with available 3D structure information. ZIKV – Zika virus (PDB ID: 5TPY)[2], MVEV – Murray valley encephalitis virus (PDB ID: 4PQV)[3], TABV – Tamana bat virus (PDB ID: 7K16)[4], POWV – Powassan virus[5], PLRV – Potato leafroll virus (PDB ID: 7JJU)[6], SCNMV – Sweet clover necrotic mosaic virus (PDB ID: 6D3P)[7].

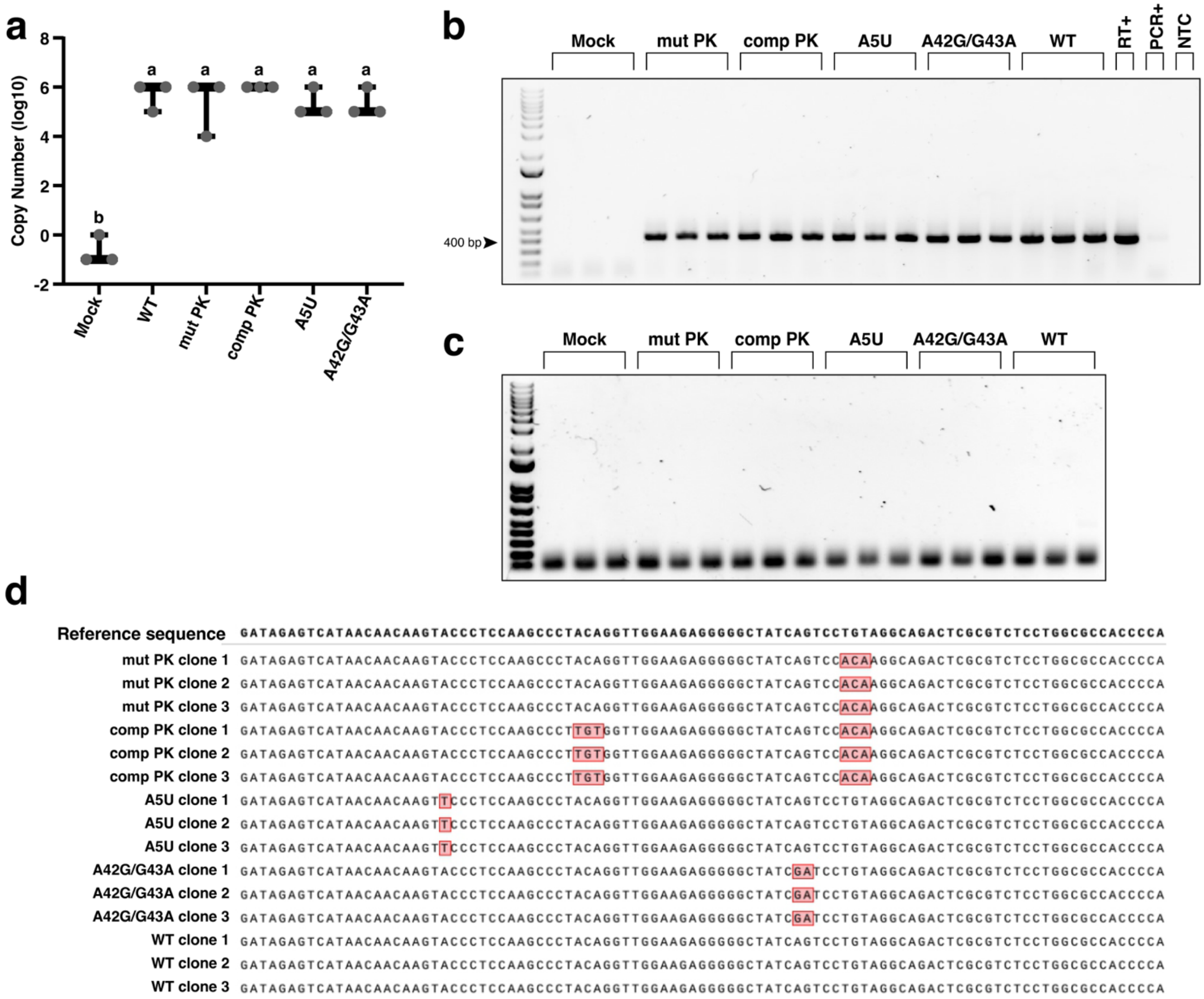


**Extended Data Fig. 2. Infection of *N. benthamiana* with ST9a.** a) Copy number of infectious ST9a RNA immediately after infiltration (time 0), quantified by qPCR. Copy number, calculated using an absolute quantification method, refers to the amount of infiltrated plasmid and is per ng of total DNA extracted from the infiltrated leaves. Data represents 3 biological replicates. "a" represents no statistical significance following One-way ANOVA, *p<0.05*, while "b" is significant. b-d) RT-PCR (b) and no RT control (c) of the indicated ST9a RNAs from agroinoculated *N. benthamiana* leaves infected for 8 days. PCR products were purified and submitted for Sanger sequencing to confirm retention of the introduced mutations (d). PCR+: WT ST9a plasmid used as a positive control; NTC: no template control.

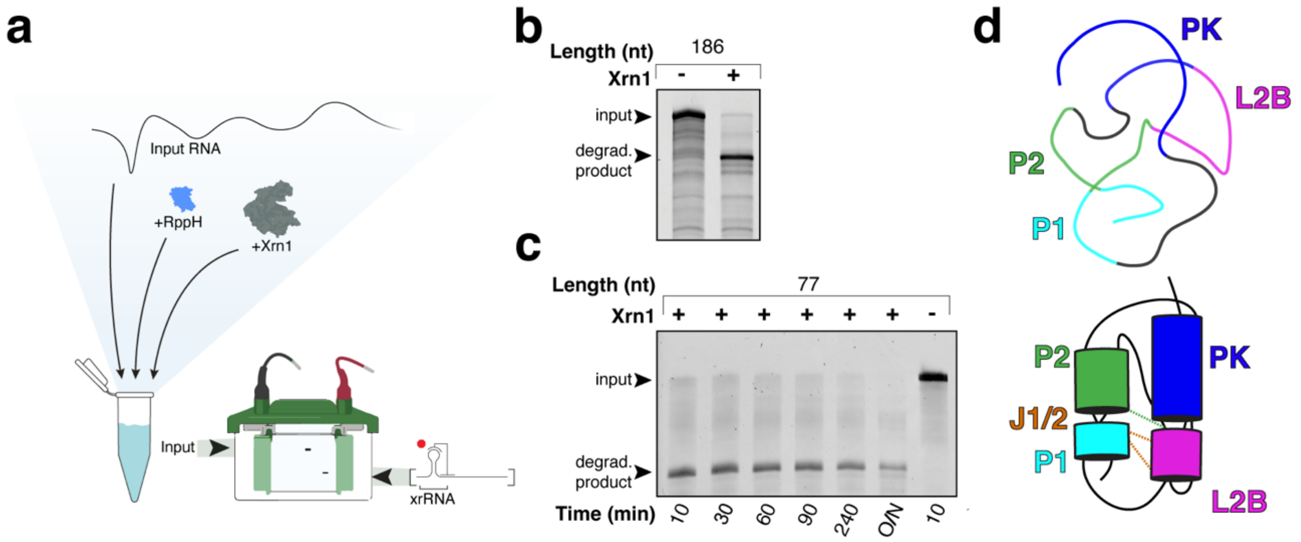


**Extended Data Fig. 3. Biochemical analyses of the ST9a xrRNA sequence.** a) Scheme of the *in vitro* Xrn1 degradation assay. b-c) *In vitro* Xrn1 degradation reaction of RNA containing the ST9a xrRNA. Reactions were resolved by dPAGE and visualized by ethidium bromide staining. Length in nucleotides (nt) corresponds to the resistant product, counting from the Xrn1 stop site as determined in Fig. 1b. Reactions were incubated for 1.5 hours (b) or the indicated times (c) at 30 ºC, resolved by dPAGE and visualized by ethidium bromide staining. O/N = overnight (~12 hours). d) Ribbon diagram (top) and cartoon model (bottom) depicting the coaxial stacking of ST9a xrRNA helices.

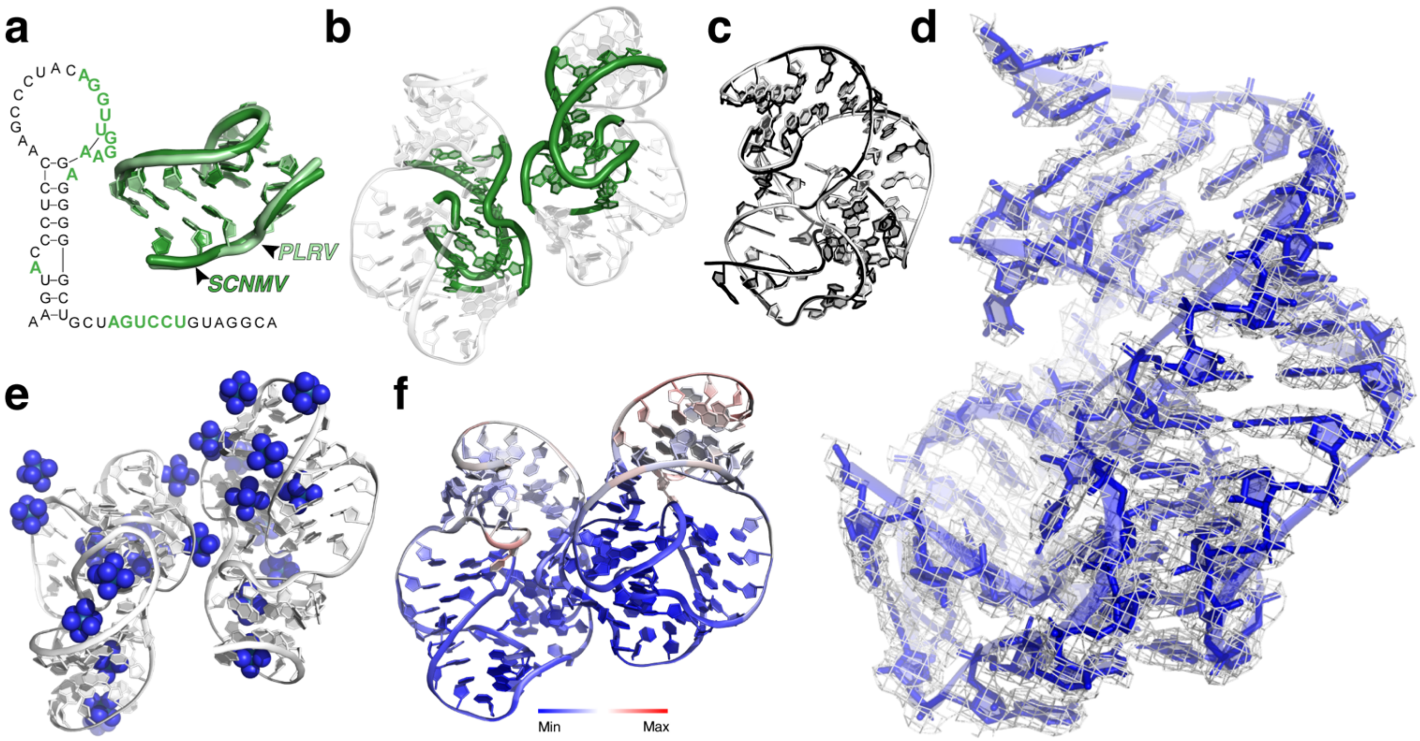


**Extended Data Fig. 4. The ST9a xrRNA crystal structure solved by a minimal molecular replacement search model.** a) The ensemble search model used for MR-SAD (right), based on an alignment of highly conserved regions of the PLRV xrRNA (PDB: 7JJU)[6] and SCNMV xrRNA (PDB: 6D3P)[7] structures, with base identities mutated to match ST9a. Left: 2D representation of the ST9a xrRNA structure with nucleotides matching the search model highlighted in green. b) MR search ensemble (in green) placed within the 2 RNA molecules of the asymmetric unit. The final structure model after refinement is shown in white. c) Overlay of the two chains of the asymmetric unit shows high agreement with an RMSD of 0.237 Å. d) Final 2Fo-Fc electron density map after model building and refinement (grey mesh, Contour Level=1.0 and Carve=1.5) superimposed on the final model. e) 24 iridium(III) hexammine ions (blue) in the asymmetric unit placed by MR-SAD. f) ST9a xrRNA structure colored according to B-factors, with the minimum B-factor at 10.8 and the maximum at 174. The highest B-factors are observed at the 3′ end and in the PK region.

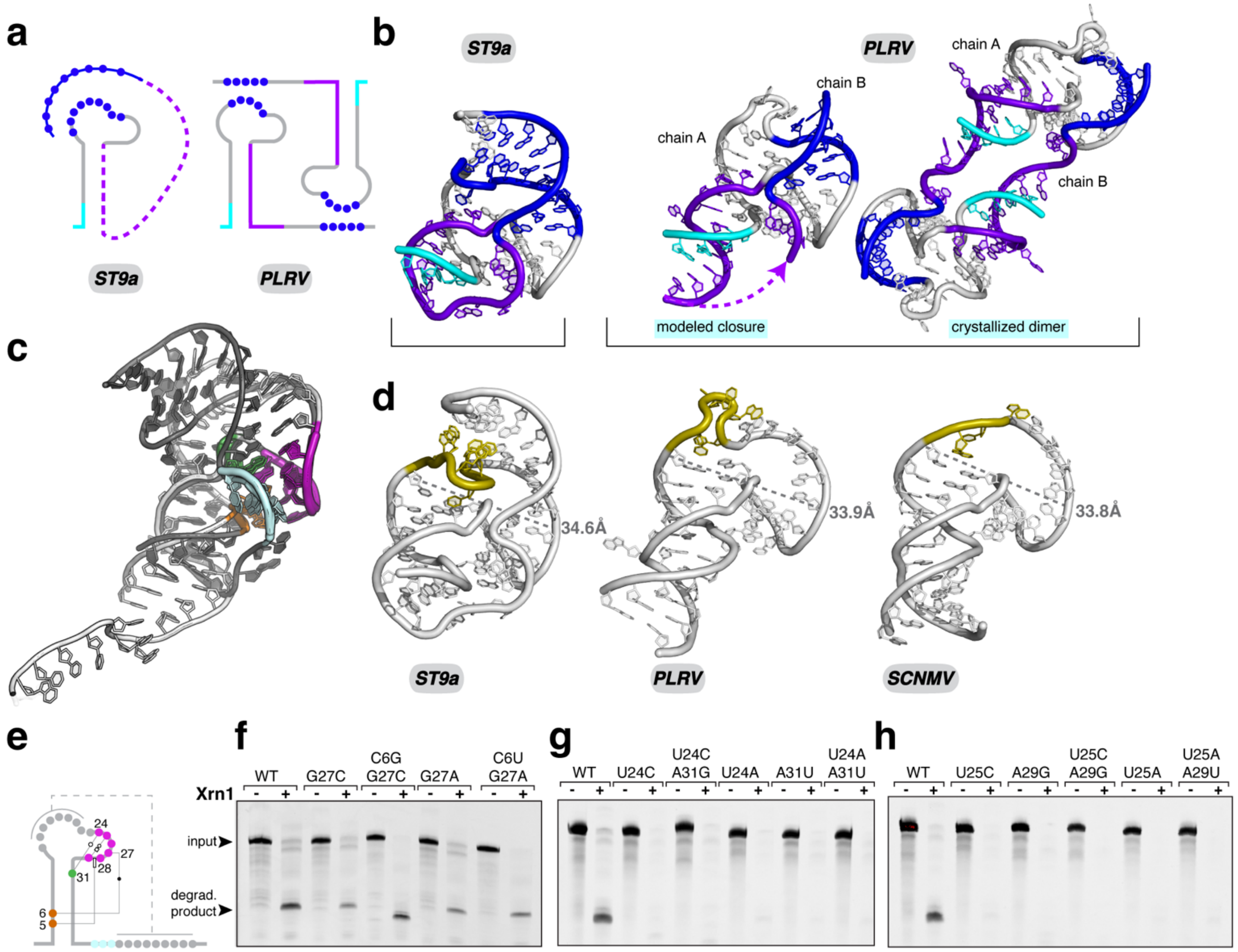


**Extended Data Fig. 5. Intramolecular contacts of the ST9a xrRNA structure form a pseudoknot.** A) Schematic representation of the PK interactions of class 3 xrRNAs *in cis* (left, based on the ST9a xrRNA structure) and *in trans* (right, based on the crystallographic dimers of PLRV and SCNMV xrRNA structures[6,7]). b) Comparison of the of the ST9a xrRNA structure (left) with PK *in cis*, to the modelled *in cis*-PK conformation of the PLRV xrRNA (center), based on the crystallized PLRV xrRNA dimer (PDB: 7JJU[6], right). Colors match (a). c) Overlay of the open SL xrRNA structure from PLRV (white, PDB: 7JJU[6]) and the closed PK xrRNA structure from ST9a (dark grey, this study). Both structures were aligned based on the L2B region (thickened backbone, with colors to match Fig. 3a. d) Comparison of the variable L2A region (yellow) of ST9a xrRNA (left), PLRV xrRNA (PDB ID: 7JJU[6], middle) and SCNMV xrRNA (PDB ID: 6D3P, right). e) Schematic representation of ST9a xrRNA with mutated long-range network highlighted. f-h) *In vitro* Xrn1 degradation reaction of ST9a xrRNA WT and the indicated mutants. Reactions were resolved by dPAGE and visualized by ethidium bromide staining. Mutations of the C6-G27 base pair weaken, but do not fully abolish, Xrn1 resistance (f).

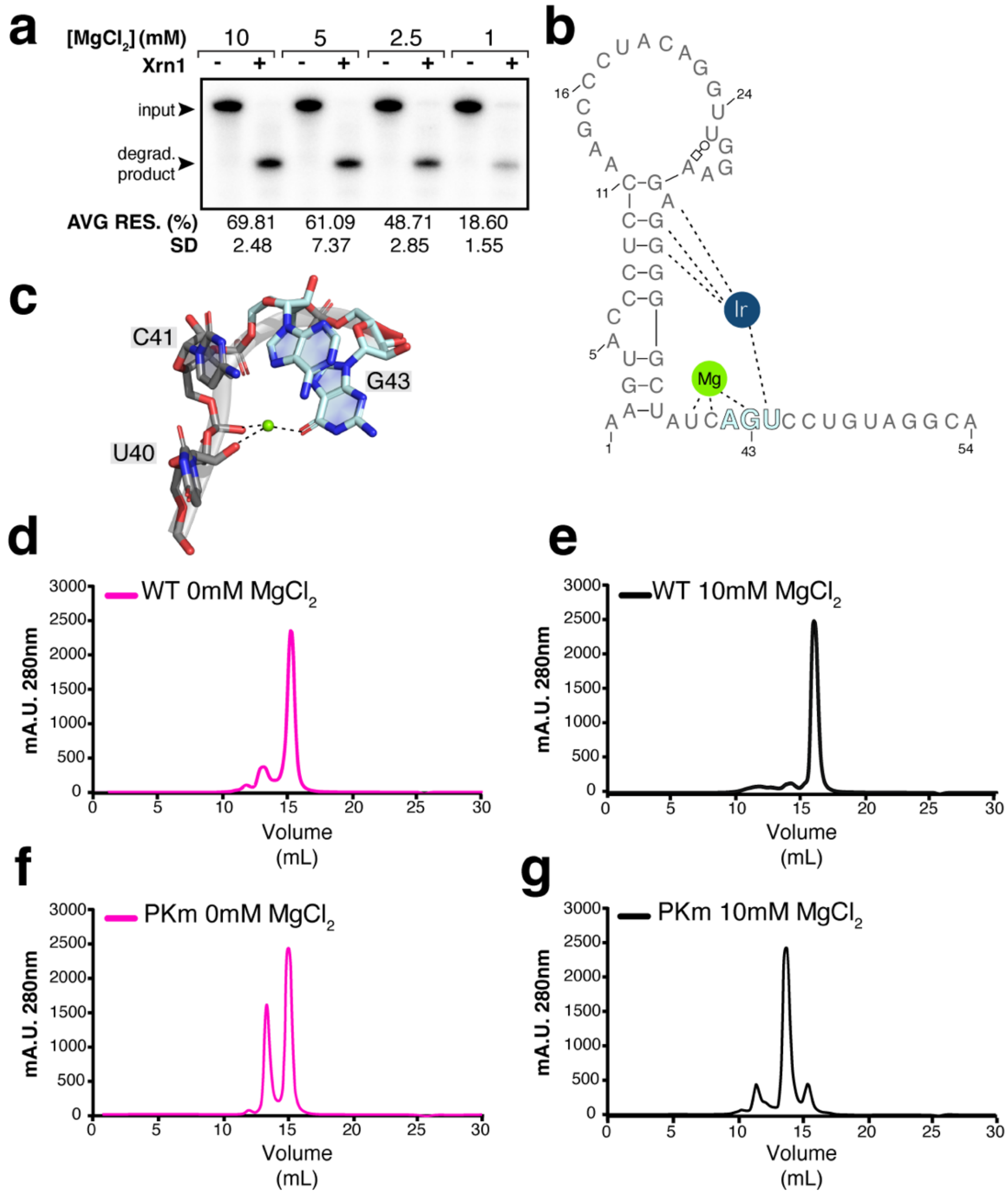


**Extended Data Fig. 6. Coordinated metal ions stabilize the xrRNA structure in solution.** a) Quantitative *in vitro* Xrn1 degradation reaction of $^{32}$P-3′-labeled ST9a xrRNA at the indicated $MgCl_2$ concentrations. Reactions were resolved by dPAGE and visualized by autoradiography. Gel quantification shown in Fig. 4a. Average resistance (AVG RES.) from n=4 technical replicates with standard deviation (SD) shown below a representative gel. b) Secondary structure diagram of ST9a xrRNA highlighting select metal ion binding sites. c) Coordination of a $Mg^{2+}$ ion by U40, C41 and G43. d-g) SEC elution profiles for ST9a WT xrRNA without $MgCl_2$ (d), ST9a WT xrRNA with 10 mM $MgCl_2$ (e), ST9a PK mutant (PKm) without $MgCl_2$ (f), and ST9a PKm with 10 mM $MgCl_2$ (g) correlating to the respective SAXS frames shown in Supplementary Fig. 1.

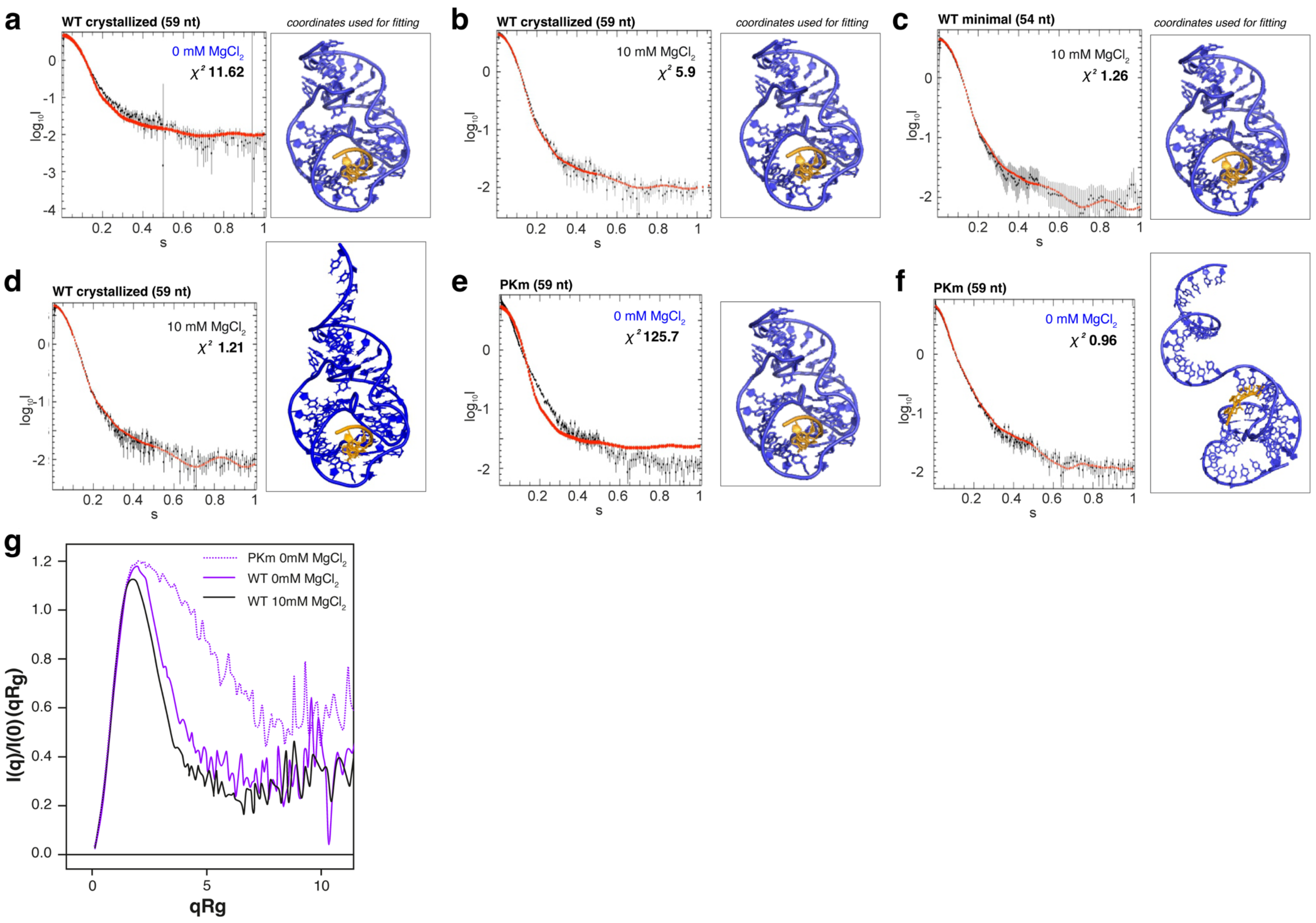

a
WT crystallized (59 nt)
coordinates used for fitting
0 mM MgCl$_2$
$\chi^2$ 11.62
b
WT crystallized (59 nt)
coordinates used for fitting
10 mM MgCl$_2$
$\chi^2$ 5.9
c
WT minimal (54 nt)
coordinates used for fitting
10 mM MgCl$_2$
$\chi^2$ 1.26
d
WT crystallized (59 nt)
10 mM MgCl$_2$
$\chi^2$ 1.21
e
PKm (59 nt)
0 mM MgCl$_2$
$\chi^2$ 125.7
f
PKm (59 nt)
0 mM MgCl$_2$
$\chi^2$ 0.96
$log_{10}I$
s
g
PKm 0mM MgCl$_2$
WT 0mM MgCl$_2$
WT 10mM MgCl$_2$
I(q)/I(0) (qRg)
qRg

**Extended Data Fig. 7. Fitting of in-solution scattering profiles with calculated curves based on the crystal structure.** SAXS raw scattering curves in presence of varying $Mg^{2+}$ concentrations for ST9a xrRNA constructs as indicated, overlaid with calculated theoretical curves (red) derived from structural model coordinates of the ST9a xrRNA (right). The fit is given by the χ2 value. a) Scattering curve for the 59-nt RNA crystallization construct in 0 mM $Mg^{2+}$ overlaid with theoretical curve derived from the 55-nt RNA structure model built by crystallography (PDB ID: 9CFN) (χ2=11.62). Residues 56-59 were not included in the final model due to insufficient electron density. b) Scattering curve for the 59-nt crystallization construct in 10 mM $Mg^{2+}$ overlaid with theoretical curve derived from the 55-nt model shows an improved fit relative to a) (χ2=5.9). c) Scattering curve for the 54-nt minimal xrRNA in 10 mM $Mg^{2+}$ overlaid with theoretical curve derived from the 55-nt model shows a good fit (χ2=1.26). d) Scattering curve for the 59-nt crystallization construct in 10 mM $Mg^{2+}$ overlaid with theoretical curve based on the 55-nt model extended by residues 56-59, built in COOT without adjustment to the density, also shows a good fit (χ2=1.21). e) Scattering curve for 59-nt ST9a xrRNA with mutated U49A/A50U (PKm) shows extremely poor fit to a curve calculated based on the ST9a xrRNA crystal structure (χ2=125.7). f) Scattering curve for the 59-nt ST9a-PKm xrRNA shows good fit with a 59-nt ST9a xrRNA modeled in an open conformation without PK with RNAmasonry[8] (χ2=0.96). g) Dimensionless Kratky plots of the monomeric SEC peak fractions of the indicated RNA samples.

```
              P1      J1/2  P2      L2A   PK        L2B       P2         J2/1  P1      J1/3         PK
➔ ST9.a       AAGU--AC-CCUCC--AAGC-CCUACAGG-UUGGAA--G--A-GGGG----GCU---AUC---AGU-CCUGUAGGCA
CABYV.a       CAGU--AC-CCCCC--AAGU-CCGAUGGG-UUGGAA--G--A-GGGG----GCU---AUC---AAC-CCCACCGGCA
TuYV_DSMZ.a   AAGU--AC-CCUCC--AAGU-CCUACGGG-UUGGAA--G--A-GAGG----GCU---AAC---AGU-CCCGUAGGCA
TuYV_Landkreis.a CAGU--AC-CCCCC--AAGU-CCGAUGGG-UUGGAA--G--A-GGGG----GCU---AUC---AAC-CCCAUUGGCA
* PLRV        AACU--AG-CCAAGC-AUAC---ACGAG-UUGCAA-GC-A-UUGG-A---AGUU-CA----AGC-CUCGU---UA
* SCNMV       GCGU--AA-CCUCC--AUC------CGAG-UUGCAA-G--A-GAGG-GAA-ACGC--------AGU-CUCG----CC
RCNMV         GUGU--AG-CCUCC--ACC------CGAG-UUGCAA-G--A-GGGG-A---ACAC--GC----AGU-CUCG----CC
CRSV          CCGU--AG-CCGCC--AA------CAAAG-UUGCAA-G--A-GCGG-----GCGUU-GCU---AGC-CUUUG---CC
MCMV          GGUG--AG-CCGGC--AU-------GAGG-UUGCAA-G--A-CCGG-AA--CAACC-------AGU-CCUU----CU
OPMV          CCAC--AG-CCAAGC-A-------UUAAG-UUGCAA-GC-G-UUGG-A---GUGGC-------AGG-CUUAA---CG
TBTV          AACGG-AG-CUAGA--UU------UAGUG-UUGCAA-UU--G-CUAG-A---CCGUU-------AGC-CACUA---CG
IYMV2         CCGU--AG-CCCGC--GUU------GUAG-UUGCAA-G--A-CGGG-A---GCGUU-------AGU-CUAC----UG
CMoMV         ACGU--AG-CUAGUA---------CCAGG-UUGCAA-U--A-CUAG-A---GCGUU-------AGU-CCUGG---GG
WLYaV         ACGUC-AG-CCGCC--A------ACACAG-UUGCAA-G--A-GCGG-AA--GACGU-------AGU-CUGUGU--CA
ScYLV         UCCCG-AG-CCACC--A------UAUAGG-UUGCAA-G--A-GUGG-AA--CGGGA-------AGU-CCUAUA--GA
SaYV          GAACC-AG-CGAAC--A------AUUCAG-UUGCGA-G--A-UUCG-A---GGUUU-C-----AGU-CUGAAU--CA
WCIMV         AGAU--AG-CCGGUA-UCA------GUUG-UUGCAA-UA-G-CCGG-A---GUCUU-------AGU-CAAC----GU
MABYV         CACU--AG-CCGGAA-AUAC-----GUUG-UUGCAA-UU-G-CCGG-A---AGUUU-A-----AGC-CAAC----UA
SABYV         UAACU-AG-CCGGAA-AUAC-----GUUG-UUGCAA-UU-G-CCGG-A---AGUUA-A-----AGU-CAAC----UA
CYDV-RPV      AACU--AG-CCGGAC-AAAC----GUAAG-UUGCAA-GU-G-CCGG-A---AGU---CA----AGU-CUUAC---AC
MYDV-RMV      AGUC--AG-CCAGGC-AAAU----UCGAG-UUGCAA-GC-A-CUGG-A---UGACCU------AGU-CUCGA---UA
LeYV          AGCC--AG-CCACAC-AUAC----ACAAG-UUGCAA-GU-A-GUGG-A---GGUU--C-----AGU-CUUGU---UA
CRLV          AGCU--AG-CCGAGAAACAC-----GUUG-UUGCAA-UC-G-UCGG-A---AGC---AA----AGC-CAAC----UA
CpPV2         AGUU--AG-CCGGAA-AGUC-----GCUG-UUGCAA-UU-G-CCGG-AA--AACU--U-----AGU-CAGC----UA
TVDV          AACU--AG-CCAAGC-AUAC----RUCAG-UUGCRA-GC-G-UUGG-A---AGUU--CA----AGU-CUGAU---UA
CABYV         CACU--AG-CCAAGC-ACAC----ACGAG-UUGCAA-GC-A-UUGG-A---AGUCU-G-----AGU-CUCGU---UA
PVYV          AACU--AG-CCAAGC-AUAC----AUCAG-UUGCAA-GC-G-UUGG-A---AGUU--CA----AGU-CUGAU---UA
PABYV         AAGCC-AG-CCGAGU-CAGA-----CUUG-UUGCAA-AC-G-UCGG-A---GGCUU-A-----AGU-CAAG----CA
BWYV          CACU--AG-CCGAGC-AAAC----GAAAG-UUGCAA-GC-A-UCGG-A---AGU---CA----AGU-CUUUC---AC
BChV          AAAC--AG-CCGGGU-AAAC----AUCAG-UUGCAA-AC-A-CCGG-AA--GUUU--U-----AGU-CUGAU---UA
BVG           AAGCU-AG-CCAGAC-AUAC----GUGAG-UUGCAA-GU-A-CUGG-A---UAGCUU------AGU-CUCAC---AC
OV5           AACAU-AG-CCUCAC-AUAC-----ACAG-UUGCAA-GU-G-GAGG-A---GUGUU-U-----AGU-CUGU----UA
BrYV          AGUU--AG-CCCUAC-AUAC----GUAAG-UUGCAA-GU-A-AGGG-A---AAC---AA----AGU-CUUAC---AC
PDMV          AGUG--AG-CCAGAC-AUAC----GCAAG-CUGCAA-GU-A-CUGG-A---CACU--A-----AGC-CUUGC---AC
PeWBVYV       AGUU--AG-CCAAGC-AUAC----AUCAG-UUGCAA-GC-G-UUGG-A---AACU--UA----AGU-CUGAU---UA
UPolV1        AGUA--AG-CCAAGC-AUAC----AUCAG-UUGCAA-GC-A-UUGG-AA--UACU--U-----AGC-CUGAU---UA
ZABYV         AGAC--AG-CCAGAC-ACAC----GCAAG-UUGCAA-GU-A-CUGG-AA--GUCU--UA----AGU-CUUGU---UA
PuPV          AGCC--AG-CCGAGU-CAGA-----CUUG-UUGCAA-AC-G-UCGG-A---GGCU--AA----AGU-CAAG----CA
PhBMYV        CACU--AG-CCAAGC-AAAC----ACGAG-UUGCAA-GC-A-UUGG-A---AGU---AAA---AGU-CUCGU---UA
TV2           AGCU--AG-CCAAGC-AUAC----ACGAG-UUGCAA-GC-A-UUGG-A---AGUU--CA----AGC-CUCGU---UA
AEYV          GAUC--AG-CCAAGC-AUAC----AUCAG-UUGCAA-GC-G-UUGG-A---GAUC--AC----AGU-CUGAU---UA
LABYV         AUAC--AG-CCGGGC-AUAC----CGUAG-UUGCAA-GC-G-CCGG-AAA-GUAU--CUG---AGU-CUACG---UA
WaPV          AGCC--AG-CCGAGU-UAGA-----CUUG-UUGCAA-AC-G-UCGG-A---GGCU--AA----AGU-CAAG----CA
ALVE          ACAC--AG-CCAAUA-------AUCUUG-UUGCAA-UA-G-UUGG-A---GUGU--UC----AGC-CAAGAU--GC
CpCSV         GAGAU-AG-CCGAC--U------AACUAG-UUGCGA--G-A-UCGG-A---GUCUC-------AGU-CUAGUU--CA
CpPV10        GAAUC-AG-CCCUAC-------AUCAAG-UUGCGA-GU-G-AGGG-A---GAUUC-------AGC-CUUGAU--UC
CoLRDV        GAGCU-AG-CUCUUA-------AUCUUG-UUGCAU-UA-G-AGAG-A---AGCUU-AA----AGU-CAAGAU--CA
ZPE1          CAGU--AG-CUGAAG-C-------GAAAG-UUGCAA-CU-G-UCAG-A---ACUG--U-----AGU-CUUUC---AC
HuPLV1        GAGUC-AG-CCAAAC-AAAC----ACAAG-UUGCAA-GU-G-UUGG-A---GACUC-AUUCU-AGU-CUUGU---UA
```

3D ring

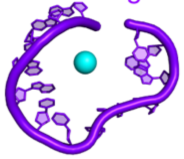

**Extended Data Fig. 8. Widespread distribution of putative class 3 xrRNAs.** Representative alignment showing class 3 xrRNA conservation. Region bracket colors correspond to colors in Fig. 1d. The ST9a sequence is indicated with a purple arrow, and the xrRNA sequences for which a crystal structure exists- PLRV (PDB: 7JJU[6]) and SCNMV (PDB: 6D3P[7]), are indicated with an asterisk.

**Supplementary Table 1.** SAXS data acquisition, sample details and data analysis for ST9a WT and PK mutant RNAs in two different buffer conditions (0 mM vs 10 mM $MgCl_2$).

| **Sample details** | **ST9a WT monomer** | | **ST9a PK mutant monomer** | | **ST9a PK mutant dimer** | |
|---|---|---|---|---|---|---|
| **$MgCl_2$ (mM)** | **0** | **10** | **0** | **10** | **0** | **10** |
| **SASBDB accession codes** | **SASDVY4** | **SASDVX4** | **SASDV25** | **-** | **SASDV35** | **SASDVZ4** |
| **Organism** | **Beet Western Yellows ST9 associated, tombusvirus-like associated RNA (tlaRNA)** | | | | | |
| **Genomic boundaries (5′ -> 3′) (GeneBank_1782103816)** | **2378-2436** | | | | | |
| **Number of nucleotides** | **59** | | | | | |
| **Mutation in sequence (GeneBank_1782103816)** | **-** | **-** | **U2326A, A2327U** | **U2326A, A2327U** | **U2326A, A2327U** | **U2326A, A2327U** |
| **Ext. coefficient ε 260nm ($M^{-1}$ $cm^{-1}$)** | **582600** | | | | | |
| **Data collection parameters** | | | | | | |
| **Instrument** | **16-ID (LiX) beam line at the National Synchrotron Light Source II (NSLS-II; Upton, NY, USA)** | | | | | |
| **Data collection mode** | **SEC-SAXS** | | | | | |
| **X-ray wavelength (nm)** | **0.082** | | | | | |
| **Sample-to-detector distance (m)** | **3.56** | | | | | |
| **Detector** | **Pilatus 1M** | | | | | |
| ***q* measurement range ($Å^{-1}$)** | **0.06-32** | | | | | |
| **SEC column** | **Superdex200 Increase 10/300** | | | | | |
| **SEC flow rate (mL $min^{-1}$)** | **0.75** | | | | | |
| **SEC-SAXS buffer** | **50 mM Tris-HCl, 100 mM natrium chloride, 1mM DTT, variable $MgCl_2$ concentration (see second line for details)** | | | | | |
| **SEC temperature (°C)** | **20** | | | | | |
| **Sample injection volume (µL)** | **95** | | | | | |
| **RNA sample conc. (mg $mL^{-1}$)** | **3** | | | | | |
| **No. of frames collected** | **1200** | | | | | |
| **Data processing** | | | | | | |
| **SEC-SAXS primary data processing** | **BioXTAS RAW (2.2.1) and CHROMIXS** | | | | | |
| **# buffer frames used for averaging** | **193** | **85** | **175** | **-** | **189** | **141** |
| **#sample frames used for averaging (frame selection)** | **13** | **13** | **15** | **-** | **23** | **12** |
| ***q* working range ($Å^{-1}$)** | **0.14-10** | **0.14-10** | **0.12-10** | **-** | **0.06-10** | **0.1-10** |
| **Data analysis** | **PRIMUS (ATSAS 4.0.0-2) and BioXTAS RAW (2.2.1)** | | | | | |
| **Structural parameters** | | | | **-** | | |
| **Guinier analysis** | | | | **-** | | |
| **Data analysis software** | **PRIMUS (ATSAS 4.0.0-2)** | | | | | |
| **$R_g$, σ (nm)** | **2.09 (+/- 0.01)** | **1.91 (+/- 0.01)** | **2.53 (+/- 0.02)** | **-** | **3.79 (+/- 0.01)** | **3.52 (+/- 0.02)** |
| **$qR_g$ limits (point range)** | **0.35-1.25 (12-50)** | **0.38-1.3 (15-54)** | **0.35-1.26 (9-45)** | **-** | **0.49-1.29 (8-29)** | **0.46-1.27 (8-31)** |
| **pI analysis** | | | | | | |
| **Data analysis software** | **RAW 2.2.1 (GNOM)** | | | | | |
| **$R_g$, σ (nm)** | **2.21 (+/- 0.02)** | **1.97 (+/- 0.007)** | **2.62 (+/- 0.01)** | **-** | **3.92 (+/- 0.04)** | **3.67 (+/- 0.01)** |
| **$D_{max}$ (nm)** | **8.8** | **7** | **9.1** | **-** | **13.5** | **13** |
| **Molecular weight** | | | | | | |

| | | | | | | |
|---|---|---|---|---|---|---|
| **Calculated MW from sequence (kDa)** | **19.23** | **19.23** | **19.23** | **19.23** | **38.46** | **38.46** |
| **MW from SAXS data (credibility range) (kDa)** | **20.6 (19.6-21.5)** | **18.7 (17.8-20.2)** | **23.1 (20.9-23.4)** | **-** | **45.7 (43.3-50.3)** | **43.8 (41.5-45.2)** |
| **Atomistic modeling** | | | | | | |
| **Structure / Model type** | **9CFN (crystal structure)** | **9CFN (crystal structure)** | **9CFN (crystal structure) / RNA Masonry** | **-** | **-** | **-** |
| **Data analysis software** | **CRYSOL (ATSAS 4.0.0-2)** | | | | | |
| ***q* range for fitting (Å$^{-1}$)** | **1.5** | | | | | |
| **Imposed symmetry** | | | **/ -** | **-** | **-** | **-** |
| **Quality of fit ($\chi^2$)** | **11.62[a]** | **5.92[a]** | **125.7 / 0.958** | **-** | **-** | **-** |

**[a]SAXS data were recorded for the 59-nt crystallization construct. However, nucleotides 56-59 were not included in the final model due to insufficient electron density. This explains the high $\chi^2$ values for experimental SAXS data fitted to the crystal structure 9CFN. See also Extended data Fig. 7.**

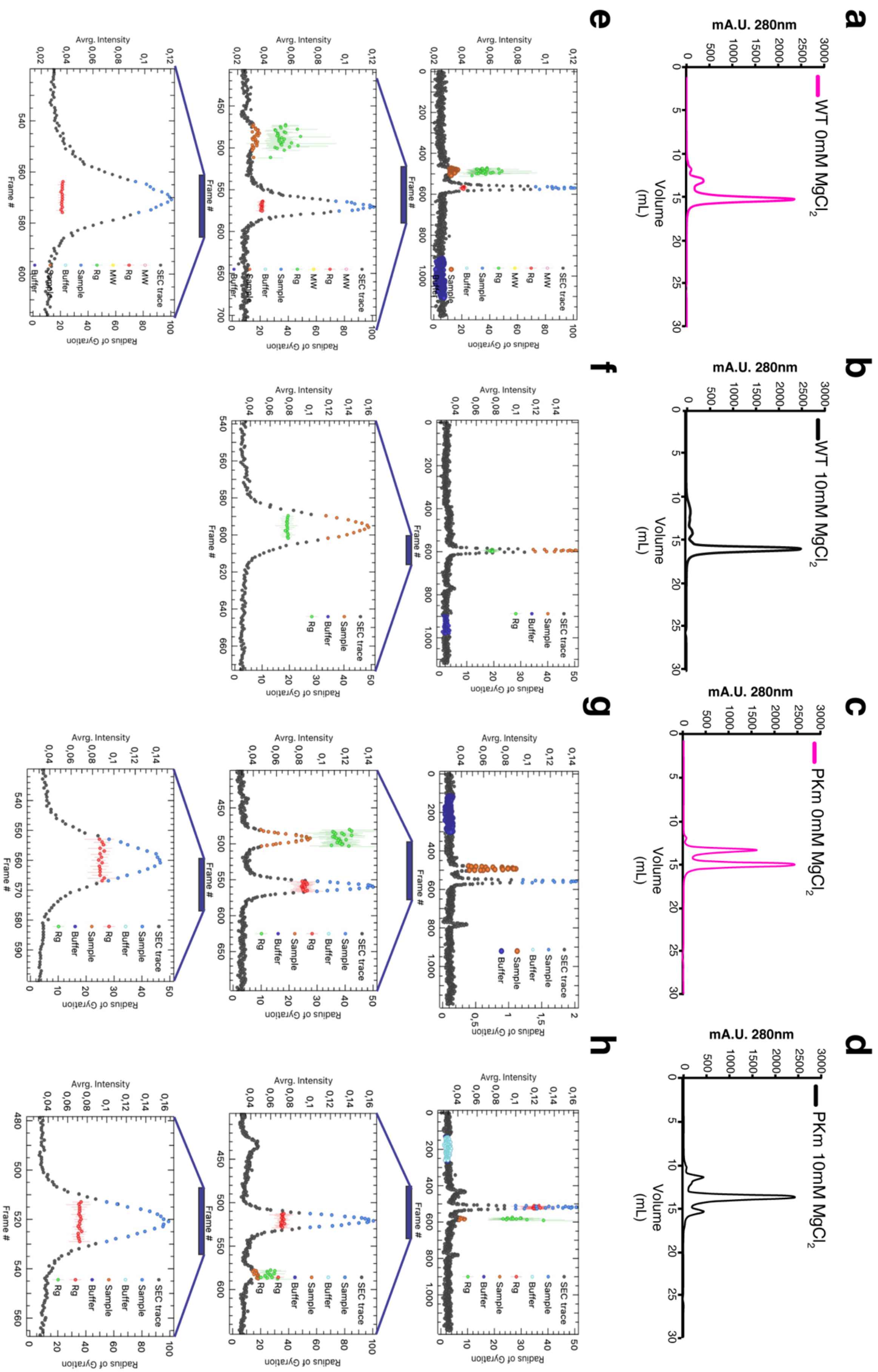

a
b
c
d
e
f
g
h
mA.U. 280nm
WT 0mM $MgCl_2$
WT 10mM $MgCl_2$
PKm 0mM $MgCl_2$
PKm 10mM $MgCl_2$
Volume (mL)
Avrg. Intensity
Frame #
Radius of Gyration
SEC trace
Sample
Buffer
Rg
MW

**Supplementary Fig. 1. SEC Frame selection for ST9a xrRNA SAXS analysis.**

a-d) SEC profiles for ST9a WT xrRNA without $MgCl_2$ (a), ST9a WT xrRNA in 10 mM $MgCl_2$ (b), ST9a PKm xrRNA without $MgCl_2$ (c), and ST9a PKm xrRNA with 10 mM $MgCl_2$ (d), with correlating SAXS frames (e, f, g, and h, respectively), shown with different zoom-in regions. Selected sample frames (blue or orange) and buffer frames (dark blue) are indicated. The *Rg* (in Å) is given for the selected sample frames (red or green). SEC profiles are recorded at 280 nm. SEC buffer was 50 mM Tris-HCl, 100 mM NaCl, 1 mM DTT, either without or with 10 mM $MgCl_2$.

### Materials and Methods

#### RNA preparation

RNA constructs were *in vitro* transcribed using T7 RNA polymerase. Transcription templates were generated by PCR amplification using custom oligonucleotides (0.5 µM each) (IDT) (see Supplementary Table 3), Phusion Hot Start DNA polymerase (New England Biolabs), and HF Buffer (Thermo Fisher Scientific). Gene fragments (Twist Biosciences) served as the PCR templates. Constructs for Xrn1 degradation assays contained the xrRNA sequence plus 60-nt endogenous upstream sequence to allow loading of the exoribonuclease. The entire PCR mix was added directly as 25% v/v to the transcription mix, which included 6 mM of each NTP, 30 mM $MgCl_2$, 30 mM Tris-HCl pH 8, 10 mM DTT, 3.5 mM spermidine, 0.1% v/v Triton X-100, 40 U/ml RNasin (Promega), and 0.05 mg/ml T7 RNA Polymerase. Transcription reactions were incubated for 4-6 hours at 37°C. Completed reactions were centrifuged for 10 min at 5000 x g to remove insoluble inorganic pyrophosphate salts. For *in vitro* Xrn1 degradation assays, the RNA products were purified using a Monarch RNA cleanup kit (New England Biolabs) according to the manufacturer's instructions. For crystallography, SAXS and thermal melting experiments, the RNA was ethanol precipitated and purified on a 12% 8 M urea Tris–borate–EDTA (TBE) polyacrylamide (19:1 acrylamide:bisacrylamide) gel. The RNA band of the correct size was excised and soaked in 0.3 M sodium acetate pH 6.7 for at least 12 hours. The liquid was filtered using 0.22 µM Steriflip vacuum filtration units (Millipore), concentrated using a 10 kDa molecular weight cutoff Amicon centrifugal concentrator (Millipore) and washed at least 10 times with nuclease-free milliQ water. Concentrated RNAs were stored at -20 ºC until use.

#### Protein Purification

The expression vector for *Kleuveromyces lactis* Xrn1 (residues 1-1245)[9] was a gift from Prof. Liang Tong at Columbia University, and the expression vector for *Bdellovibrio bacteriovorus* 5′-RNA Pyrophosphohydrolase (RppH)[10] was a gift from Joel Belasco at New York University. All recombinant proteins were 6xHis-tagged, expressed in *E. coli* LOBSTR cells and purified using Ni-NTA resin (Thermo Fisher Scientific). The protein was dialyzed into storage buffer containing 20 mM Tris pH 8, 300 mM NaCl, 2mM BME, and 10% glycerol and stored at -80 ºC. The purity of the recombinant proteins was verified by stain-free SDS-PAGE (Bio-Rad).

#### Exoribonuclease degradation assay

4 µg of RNA was resuspended in 40 µl reaction buffer containing 100 mM NaCl, 50 mM Tris-HCl pH 7.5, 1 mM DTT, and 10 mM $MgCl_2$ (or as noted). The RNA was refolded by heating to 90°C for 3 minutes, followed by 5 min incubation at 20°C. 1.5 µg of RppH was added per sample, which was then split into two 20 µl reactions, to which we added either 0.8 µg Xrn1 (Xrn1+) or 1 µl water (Xrn1-). The reaction was incubated for 1.5 hours (or as indicated) at 30°C, analyzed by dPAGE and visualized by ethidium bromide staining.

**Quantitative exoribonuclease degradation assay**

To quantify Xrn1 resistance activity, RNAs were radiolabeled at their 3′ terminus using [5′-$^{32}$P]cytidine 3′,5′-bis(phosphate) ($^{32}$PCp). First, cytidine-3′-monophosphate (Cp) was 5′-labeled in a 10 µl reaction containing 5 µL [γ-$^{32}$P]ATP (1mCi) (Perkin Elmer), 2µL Cp (1mM), 1 µL T4 Polynucleotide Kinase (PNK) (New England Biolabs), and 1 µL PNK Buffer (New England Biolabs). The reaction was incubated for 2 hours at 37 ºC, followed by 20 minutes at 60 ºC. 1 µL of the $^{32}$PCp was ligated to 5 µg Monarch-purified RNA in a 10 µL reaction volume containing 1 µL T4 RNA Ligase (New England Biolabs) and 1 µL Ligase buffer. The ligation reaction was incubated overnight at 16 ºC and purified using a P-30 spin column (Bio-Rad) according to the manufacturer's instructions. Counts per minute (cpm) were determined using liquid scintillation counting (Perkin Elmer Tri2810R). The equivalent of 25,000 cpm of $^{32}$P-3′-labeled RNA was added to a 40 µL Xrn1 degradation reaction, split into 20 µL per condition as described above. After 30 minutes incubation at 30 ºC, one fourth of each reaction (~3000 CPM per lane) was loaded onto a dPAGE gel (12% polyacrylamide) and visualized using a phosphor screen (Molecular Dynamics) and Typhoon biomolecular imager (Cytiva). Bands were quantified using ImageJ software, and Xrn1 resistance was calculated as percentage of Xrn1-resistant degradation products (resistant RNA) relative to the total input (untreated RNA). To normalize for unequal Xrn1 loading in each condition, any leftover input RNA in the treated (+Xrn1) lane was subtracted from the total input (-Xrn1) value. Four technical replicates were run in parallel and used for analysis (ordinary One-way ANOVA, $p<0.05$ and Tukey's multiple comparisons test (Prism 10).

**X-ray crystallography**

RNA was prepared as described above at a 10 mL scale. To avoid heterogeneity of 5′ and 3′ ends, the crystallization construct was flanked by a 5′ Hammerhead (HH) ribozyme and 3′ Hepatitis Delta Virus (HDV) ribozyme sequence. To activate ribozyme cleavage at the end of the transcription incubation time, the $MgCl_2$ concentration was increased to 60 mM and the reaction was incubated for 10 minutes at 65°C. The RNA was purified using dPAGE as described above.

Below is the DNA sequence used for transcription, with the ribozyme sequences underlined, the **T7 promoter in bold**, and the ***crystallization construct bolded and italicized*** (see Supplementary Table 3 for sequence information):

5′-GATCGGATCC**TAATACGACTCACTATAGGG**AAGAATGAAACTGTCACCATTTGTATAATATTTCTAAGAGGAGGGTACTTCTGATGAGTCCGTGAGGACGAAACGGTACCCGGTACCGTC***AAGTACCCTCCAAGCCCTACAGGTTGGAAGAGGGGGCTATCAGTCCTGTAGGCAGACTC***GGGCGGCATGGTCCCAGCCTCCTCGCTGGCGCCGCCTGGGCAACATGCTTCGGCATGGCGAATGGGACCCTGTTTGTTTACAATACCAGGTCAACCGCCC-3′

To screen crystallization conditions, RNA at a concentration of 5 µg/µL (~250 µM) was refolded in 2.5 mM $MgCl_2$ and 10 mM HEPES-KOH pH 7.5 by heating to 65°C for 3 minutes and cooling to room temperature before adding spermidine to a final concentration of 0.5 mM. Crystal Screens I and II, Natrix I and II, and Nucleic Acid Mini Screen (all from Hampton Research) were used to perform initial screens at 20 °C using sitting-drop vapor diffusion crystallization of 150 nL RNA solution mixed with 150 nL of reservoir solution. The RNA used for the final structural analysis was crystallized in 0.02 M magnesium chloride hexahydrate, 0.05 M sodium cacodylate trihydrate pH 7.0, 15% v/v 2-Propanol, 0.001 M hexammine cobalt(III) chloride, and 0.001 M spermine. Crystals were soaked in freezing solution – mother liquor containing 10 mM iridium(III) hexammine trichloride and 30% 2-Methyl-2,4-pentanediol (MPD) – and frozen in liquid nitrogen. Data were collected at the NYX beamline at Brookhaven National Laboratory at a wavelength of 0.9795 Å, near the Ir $L_{II}$ edge at 100K. X-ray diffraction data were indexed, integrated, and scaled using XDS[11]. Experimental phasing by single-wavelength anomalous dispersion (SAD) was not successful, so a combined molecular replacement (MR)-SAD strategy was used. The MR ensemble was based on the conserved L2B region of SCNMV and PLRV xrRNA structures (PDB 6D3P:{7, 18-26, 28 of chain A; 40-45 of chain B} and PDB 7JJU {7, 21-29, 32 of chain A; 44-49 of chain B}) with base identities mutated to match the ST9a xrRNA. The initial MR run resulted in a Translation Function Z-score (TFZ)=12.7 and Log Likelihood Gain (LLG)=221.86, and MR-SAD identified 24 iridium(III) hexammine sites. The map was used to build an initial model, which was improved through iterative rounds of model building and refinement (simulated annealing, Rigid-body, B-factor refinement) using COOT[12] and Phenix[13]. The improved model was again used as input coordinates for MR-SAD with Phenix Phaser-EP, resulting in a figure of merit=0.749 and LLG=1998.4, and the resulting map led to the final model containing nucleotides 1-55 of chain A and 2-55 of chain B. Residues 56-59 of both chain A and B, and residue 1 of chain B were not included in the final model due to insufficient electron density. Only data to 2.9 Å were used for refinement. Crystal diffraction data, phasing, and refinement statistics are listed in Table 1.

**Bioinformatics**

An initial alignment was generated from 58 previously published class 3 xrRNA sequences from plant viruses[6], the ST9a minimal xrRNA sequence, and 19 additional sequences from plant viruses obtained using the ST9a sequence as a query in a BLASTn search[14]. These sequences were manually aligned with Jalview v2.11.3.3[15], using the ST9a xrRNA secondary structure as a reference. We used Infernal v1.1.4 with the parameter -T 0 to search a database of reference viral genomes, which comprised all available viral nucleotides from green plant hosts (taxid: 33090), retrieved from National Center for Biotechnology Information (NCBI) on 23rd April, 2024. Hits from Infernal searches were manually included in the comparative sequence alignment if they fulfilled the following conditions: Infernal E value <= 10, the presence of pseudoknot, and the length of the sequence >= 40 nt. When determining the presence of pseudoknot, we set the minimal pseudoknot base-pairing length to be 4 nt, and then extended the sequence if there was a potential for forming a pseudoknot with the next 40 downstream nucleotides. The final sequence alignment included 363 unique sequences from the initial alignment and infernal hits. For the covariance model, we performed statistical validation with R-scape v2.0.4[17] and visualized it with R2R v1.0.6[19] .

Family-level taxonomy information for each sequence was obtained from National Center for Biotechnology Information (NCBI) using the nucleotide accession number. If family-level data were unavailable, we used the most specific taxonomic level that could be retrieved. Metagene information for each sequence was obtained from National Center for Biotechnology Information (NCBI) using the nucleotide accession number and the sequence coordinates. Regions upstream of the first annotated CDS or downstream of the last annotated CDS in the nucleotides were considered as 5′ UTR or 3′ UTR, respectively. Sequences that came from nucleotides without any available feature annotations were labeled as “unknown”.

**Reverse transcription of Xrn1 degradation products and mapping of the Xrn1 stop site**

To determine the Xrn1 stop site at single-nucleotide resolution, 200 µg of *in vitro*-transcribed RNA containing the ST9a xrRNA and 60 endogenous bases upstream of the presumed Xrn1 stop site was degraded using recombinant RppH (7.5 µg) and Xrn1 (4.5 µg) in 100 µL buffered conditions as described above .The degradation product was purified by phenol-chloroform extraction, ethanol-precipitated, washed once with 70% ethanol, and resuspended to 6 µg/µL in nuclease-free water. Successful degradation was verified by dPAGE. A DNA primer for reverse transcription (see Supplementary Table 3) was 5′-$^{32}$P labeled in a 20 µl reaction volume containing 70 µM DNA oligo, 2 µL [γ-$^{32}$P]ATP (1mCi) (Perkin Elmer), 2 µL PNK (New England Biolabs) and 2 µL PNK buffer for 1 hour at 37 ºC, followed by removal of unincorporated nucleotides using a P-30 spin column (Bio-Rad) and diluted in 500 µl nuclease-free water. 100 ng of RNA was mixed with 2 µl of the $^{32}$P-labeled primer in a final volume of 10 µl and refolded by heating for 2 minutes at 85 ºC followed by 5 minutes at 20 ºC and cooling to 4 ºC. The RNA was reverse transcribed by adding 1 µl dNTP mix (10 mM), 0.5 µl Superscript II (Thermo Fisher Scientific), 1 µl DTT (100 mM), 4 µl SS II buffer and 2.5 µl water and incubated for 1 hour at 42 ºC, followed by 15 minutes at 70 ºC. To hydrolyze the RNA template after reverse transcription, 2 µl NaOH (1M) was added and the reaction mix incubated for 5 minutes at 90°C, followed by cooling on ice for 1 min. 20 µl 2x RNA loading dye (New England Biolabs) was added, and 3 µl of the reaction were analyzed on a 12% dPAGE sequencing gel. The gel was run at 40W for 4 hours and visualized using a phosphor storage screen and Typhoon imager. As size controls, 100 ng of undigested RNA and 100 ng of the crystallization construct were reverse transcribed alongside the degraded RNA. A Sanger sequencing dideoxynucleotide (ddNTP) ladder of the undigested RNA was analyzed alongside the degradation product as reference for band annotation. For each nucleotide of the ddNTP ladder, 100 ng of the undigested (-Xrn1) RNA was reverse transcribed as described above with 0.5 mM of the ddNTP and the respective dNTP at a reduced concentration of 0.05 mM.

**Small-angle X-ray Scattering**

SEC-SAXS measurements (I(q) vs. q, where q is defined in equation (1) and 2θ is the scattering angle and λ the X-ray wavelength) were performed at the 16-ID (LiX) National Synchrotron Light Source II (NSLS-II) equipped with a Pilatus 1M detector, using the beamline’s standard SAXS configuration as described by Yang et al.[20]. Details are further listed in Supplementary Table 1.

$$(1)\ q = 4\pi \sin\theta / \lambda$$

RNAs were buffer-exchanged into SAXS buffer (50 mM Tris-HCl pH 7.5, 100 mM sodium chloride, 1 mM DTT, either with or without 10 mM $MgCl_2$) using 3 kDa molecular weight cutoff Amicon centrifugal concentrators (Millipore). Samples were adjusted to a final concentration of 3 mg/mL and snap frozen by incubation for 5 min at 95 °C and subsequent cooling in an ice bath. Samples were shock frozen in liquid nitrogen, transferred to the beamline, de-frozen and subjected to SEC-SAXS. 95 µL of RNA samples were loaded onto a Superdex 200 Increase 10/300GL (GE Healthcare) pre-equilibrated in SAXS buffer with or without $MgCl_2$ using an Agilent HPLC system at a flow rate of 0.75 mL/min. 1200 successive 2D SAXS data frames were collected from the continuously flowing eluate.

Data averaging, processing and visualization were done with BIOXTAS RAW suite[21] and the ATSAS 4.0.0-2 software suite[22]. We used CHROMIXS[23] for the selection of sample and buffer frames and to derive averaged SAXS data to produce final 1D profiles used for further analysis (Extended Data Fig. 7 and Supplementary Table 1). All final curves were processed to derive $R_g$, $P(r)$ and molecular weight estimates using Bayesian inference[24]. Structural parameters are reported in Supplementary Table 1, guided by the recommendations of Trewhella *et al.*[25]. Fitting of experimental data to structural models was carried out with CRYSOL[26]. We first fitted the SAXS data of 59-base ST9a WT xrRNA – the construct used for crystallization – to the final structure coordinates containing 55 of the 59 bases; but, to have a better size comparison, we also fitted the experimental data to an expanded model of ST9a that was generated using the "add residue" feature in Coot, avoiding adjustments based on density to prevent any bias (Extended Data Fig. 7d). We further used the processed SEC-SAXS data of ST9a PKm monomer (0 mM $MgCl_2$) to create a structural model with RNAmasonry[8] for the opened conformation via 50 iterative steps using CRYSOL[26,27] as a model fit procedure and manual input of secondary structure for the initial step. All SAXS data and relevant models are available in the Small Angle Scattering Biological Databank[28].

**Melting temperature analysis**

Circular dichroism (CD)-observed melting of ST9a xrRNA (ST9a WT minimal resistant element, 54-nt) was carried out on a ChirascanTM V100 CD spectrometer (Applied Photophysics, Inc., UK). RNA samples (200 µL of 30 µM) in 25 mM potassium phosphate (KPi) pH 6.5, 50 mM potassium chloride (KCl), either with or without 10 mM $MgCl_2$, were folded by 3 min denaturing (95 °C) and snap cooling on ice. Measurements were performed in a cuvette (Helma QS) with a sample length of 1 mm. Both CD and UV absorbance were monitored. Melting experiments were recorded at 266 nm from <8 to 95 °C with a heating rate of 1 °C/min. Melting temperatures were obtained by fitting normalized raw CD data with a one- or two-transition model in OriginPro (see equation 2, where A1 is the bottom and A2 is the top asymptote, logx0 the center and p the hill slope and equation 3, where A1 is the bottom and A2 is the top asymptote, logx01 the first and logx02 the second EC50, h1 is slope 1 and h2 slope 2, and p the proportion).

(2) y = y=A1+A2-A11+10(logx0-x)p

(3) y = y=A1+A2-A1p1+10logx01-xh1+1-p1+10logx02-xh2

The first derivative of the normalized CD data at 266 nm over temperature was plotted to derive transition points.

### Construction of ST9a mutants for agroinfection experiments

Mutants were introduced into a previously developed wild-type (WT) ST9a infectious clone (JL89:ST9a) to construct four ST9a mutant clones: mut PK (T47A/G48C/T49A), comp PK (T47A/G48C/T49A; A19T/C20G/A21U), A5U and A42G/G43. Mutations were introduced by PCR using the PrimeSTAR GXL Premix (Takara Bio), JL89:ST9a as template and DNA primers specified in Supplementary Table 3. Mutation-containing PCR products were assembled into the WT ST9a clone using the NEBuilder HiFi DNA Assembly kit (New England Biolabs) according to the manufacturer's protocol. Successful mutagenesis was confirmed by Sanger sequencing. The four ST9a mutants were then transformed into *Agrobacterium tumefaciens* (now known as Rhizobium radiobacter) strain GV3101, and used for the *Agrobacterium* infiltration assays.

### Agroinoculation and northern blot analysis

Cultures of *Agrobacterium tumefaciens* strain GV3101 harboring wild type or mutant ST9a constructs were cultured and resuspended to an A600 of 0.8[30], then syringe infiltrated into young *Nicotiana benthamiana* plants at the 4-6 leaf stage, using three biological replicates per treatment. At 8 days post infiltration, total RNA was extracted using TRIzol™ Reagent (Invitrogen) according to the manufacturer's instructions. Northern blot analysis was performed using 2 µg of total RNA extracts, according to a previously published protocol[31]. Hybridization with an ST9a specific RNA probe (see Supplementary Table 3) targeting the 3′ UTR was performed at 65 ºC overnight. The northern blot was visualized by autoradiography using a phosphor storage screen with exposures taken at 6 and 16 hours.

### qPCR quantification of inoculum

Immediately after agroinoculation (time 0), infiltrated leaf tissue was collected and total DNA was extracted using the Dellaporta method as previously described[32]. The extracted DNA was used as template in a 20 µL qPCR mix containing 10 µL of iTaq Universal Probes Supermix (Bio-Rad), 0.04 µL of each primer (100 µM), 0.02 µL of each probe (100 µM), and 1 µL of DNA. The thermocycling conditions were as follows: 95 ºC for 2 min, followed by 40 cycles of 95 ºC for 15 s and 60 ºC for 55 s. A 10-fold serial dilution of plasmid containing the full length WT ST9a sequences was used to generate a standard curve. The cytochrome c oxidase (cox) gene served as an internal control. Three biological replicates and two technical replicates were used for each treatment. The $\log_{10}$ copy number of the inoculum was calculated using the absolute quantification method in excel, and the results were statistically analyzed (One-way ANOVA, $p<0.05$) and graphed using GraphPad Prism 10 software. Refer to Supplementary Table 3 for primer and probe sequence information.

### RT-PCR to confirm mutations were retained

The same total RNA used for northern blot analysis was treated with RQ1 RNase-Free DNase (Promega), then used as template for reverse transcription using the SuperScript IV Reverse Transcriptase (Invitrogen) and 250 ng of random primers (Invitrogen). 1 µL of cDNA was used as template in a 25 µL PCR mix containing 5 µL of 5x Green GoTaq Flexi Buffer, 1.5 µL $MgCl_2$ (25 mM), 1 µL of each primer (10 µM), and 0.125 µL of GoTaq Flexi

DNA Polymerase. The thermocycling conditions were as follows: 95 ºC for 2 minutes, followed by 35 cycles of 95 ºC for 30 s, 50 ºC for 30 s, 72 ºC for 1 min, and a final extension at 72 ºC for 5 min. Primers were the same used to make the northern blot probe (see Supplementary Table 3). PCR products were visualized on a 1% agarose gel, purified using the NucleoSpin Gel and PCR Clean-Up Kit (Takara), and submitted for Sanger sequencing.

**Data availability**

The ST9a xrRNA structure model and data can be found in the Protein Data Bank (PDB) under accession 9CFN. All SEC-SAXS data generated in this study have been deposited in the SASBDB under the following accession codes: SASDVY4 (ST9a WT, monomer, 0 mM $MgCl_2$), SASDVX4 (ST9a WT, monomer, 10 mM $MgCl_2$), SASDV25 (ST9a PK mutant, monomer, 0 mM $MgCl_2$), SASDV35 (ST9a PK, dimer, 0 mM $MgCl_2$), SASDVZ4 (ST9a PK, dimer, 10 mM $MgCl_2$), SASDVV4 (ST9a WT 54 nucleotides, monomer, 10 mM $MgCl_2$) and SASDVW4 (ST9a WT 54 nucleotides, monomer, 1 mM $MgCl_2$).
